# miR-221/222 drive synovial fibroblast expansion and pathogenesis of TNF-mediated arthritis

**DOI:** 10.1101/2022.07.22.500939

**Authors:** Fani Roumelioti, Christos Tzaferis, Dimitris Konstantopoulos, Dimitra Papadopoulou, Alejandro Prados, Maria Sakkou, Anastasios Liakos, Panagiotis Chouvardas, Theodore Meletakos, Yiannis Pandis, Niki Karagianni, Maria Denis, Maria Fousteri, Marietta Armaka, George Kollias

**Affiliations:** Institute for Bioinnovation, Biomedical Sciences Research Centre (B.S.R.C.) “Alexander Fleming”, Vari, 16672, Greece; Department of Pathophysiology, Medical School, National and Kapodistrian University of Athens, Athens, 11527, Greece; Department of Physiology, Medical School, National and Kapodistrian University of Athens, Athens, 11527, Greece; Center of New Biotechnologies & Precision Medicine, National and Kapodistrian University of Athens Medical School, Athens, Greece; Institute for Fundamental Biomedical Research, Biomedical Sciences Research Center “Alexander Fleming”, Vari, Greece; Biomedcode Hellas SA, Vari, 16672, Greece

**Author notes:** Correspondence to: George Kollias, Biomedical Sciences Research Center (BSRC), ‘Alexander Fleming’, 34 Alexander Fleming Street, Vari, 16672 Greece, (contact details). Institute for Research in Biomedicine (IRB Barcelona), Baldiri i Reixac 10, 08028 Barcelona, Spain. Department for Biomedical Research, University of Bern, Bern, Switzerland (Panagiotis Chouvardas).

## Abstract

MicroRNAs (miRNAs) constitute fine tuners of gene expression and are implicated in a variety of diseases spanning from inflammation to cancer. miRNA expression is deregulated in rheumatoid arthritis (RA), however, their specific role in key arthritogenic cells such as the synovial fibroblast (SF) remains elusive. We have shown in the past that the expression of the miR-221/222 cluster is upregulated in RA SFs. Here, we demonstrate that miR-221/222 activation is downstream of major inflammatory cytokines, such as TNF and IL-1β, which promote miR-221/222 expression independently. miR-221/222 expression in SFs from the *huTNFtg* mouse model of arthritis correlates with disease progression. Targeted transgenic overexpression of miR-221/222 in SFs of the *huTNFtg* mouse model led to further expansion of synovial fibroblasts and disease exacerbation. miR-221/222 overexpression altered the transcriptional profile of SFs igniting pathways involved in cell cycle progression and ECM regulation. Validated targets of miR-221/222 included p27 and p57 cell cycle inhibitors, as well as Smarca1 (a chromatin remodeling component). In contrast, complete genetic ablation of miR-221/222 in arthritic mice led to decreased proliferation of fibroblasts, reduced synovial expansion and attenuated disease. scATAC-seq data analysis revealed increased miR-221/222 gene activity in the pathogenic and activated clusters of the intermediate and lining compartment. Taken together, our results establish an SF-specific pathogenic role of the miR-221/222 cluster in arthritis and suggest that its therapeutic targeting in specific subpopulations should inform the design of novel fibroblast-targeted therapies for human disease.

## Introduction

Rheumatoid arthritis (RA) is a chronic, painful and destructive disease with sustained inflammation in the joints, that eventually leads to destruction of cartilage and bone. Articular inflammation is fueled by the aberrant production of inflammatory mediators by immune and mesenchymal resident cells in the joints^1,2^. Genetically modified mouse models with deregulated expression of TNF develop an erosive polyarthritis resembling RA (*huTNFtg*)^3^ and SpAs (Spondyloarthropathies) (*TNF*^*ΔARE*4^, *TgA86*^5,6^), showing also extraarticular manifestations, such as heart valve disease and intestinal inflammation, recapitulating comorbid pathologies often observed in humans^4,7^.

Early studies in our lab established that TNF targeting of TNFR1 on synovial fibroblasts is required for the development of arthritis and suffices to orchestrate full pathogenesis^8,9^. Synovial fibroblasts (SFs – resident mesenchymal cells in the joints) in RA diverge from their physiological role to nourish and protect the joints, and undergo arthritogenic transformation. Under chronic inflammatory signals, SFs hyperproliferate, express ECM degrading enzymes and acquire an aggressive, invasive and tissue destructive phenotype^10–13^. In addition, arthritogenic SFs can migrate via the circulation to distant sites and transfer disease in both mice and humans^14,15^. Epigenetic alterations, such as global DNA hypomethylation and deregulated expression of microRNAs and lncRNAs^16–20^, have been linked to the aggressive and destructive behavior of SFs.

Altered expression and important functions have been reported for a number of microRNAs in RA^18,19^. microRNAs are small regulatory RNAs (19-25nt long), that fine-tune the expression of several target mRNAs and have fundamental roles in development, physiology and disease. They recognize their RNA targets by binding mainly to their 3’UTR region, guide them to the RNA-induced silencing complex (RISC) and lead them either to target degradation or translational repression^21,22^.

Specifically, miRNAs miR-221 and miR-222 have been found upregulated in SFs derived from the *huTNFtg* mouse model and RA patients^23^. miR-221 and miR-222 are clustered on the X chromosome and classified as oncomiRs, as they are upregulated in a number of human epithelial cancers. They target genes mainly involved in cell proliferation and survival, such as cell cycle inhibitors p27 and p57^24–27^. Under non-resolving inflammatory signals, such as in sepsis, the overexpression of miR-222 contributes to morbidity by targeting Smarca4, a chromatin remodeling component that participates in the SWI/SNF complex^28^. Of note, a recent study highlighted a protective role for miR-221/222 in DSS colitis in T cells by downregulating IL-23 signaling^29^. In RA, expression levels of these two miRNAs positively correlate with disease progression^30^ and *ex vivo* downregulation of miR-221 has been shown to decrease migration and invasion of human RA SFs^31^. However, the *in vivo* role of these two microRNAs in arthritis remained unexplored.

Here, we show that both TNF and IL-1β upregulate miR-221 and miR-222 expression in SFs. Transgenic mice (*TgColVI-miR-221/222*) overexpressing these two miRNAs in cells of mesenchymal origin, including SFs, when crossed with the arthritic *huTNFtg mice* develop more aggressive disease, associated with enhanced expansion of SF populations. RNA profiling of SFs from these mice revealed activated pathways involved in cell proliferation and repressed pathways associated with extracellular matrix (ECM) remodeling. Additionally, bioinformatic target prediction tools for miR-221/222 combined with RNA expression data, uncovered both known and novel candidate targets for these two miRNAs, such as cell cycle inhibitors p27 and p57, as well as the chromatin remodeling component Smarca1. Furthermore, genetic ablation of miR-221/222 in the *huTNFtg* model led to amelioration of arthritis, decreased expansion of fibroblasts and partial de-repression of the expression levels of their targets. scATAC-seq analysis of SFs from arthritic mice revealed increased miR-221/222 gene activity in the destructive subpopulations of the lining and expanding intermediate compartment. Finally, transcription factor motif enrichment analysis coupled with integrative analysis of scATAC-seq and scRNA-seq, uncovered known (NF-Kb, AP-1^32,33^) and novel transcription factors [Nrf2 (Nfe2l2), Bach1], that could potentially regulate the expression of those two microRNAs.

Overall, we provide evidence that miR-221/222 play a key pathogenic role in TNF-driven arthritis and suggest that therapeutic targeting of these two miRNAs in specific subpopulations could be beneficial for the treatment of human arthritis and perhaps other inflammatory diseases associated with pathogenic expansion of fibroblasts.

## Results

### miR-221 and miR-222 induction in *huTNFtg* SFs depends on the TNF/TNFR1 axis and IL-1β

We have previously shown that miR-221 and miR-222 levels are increased in SFs from arthritic 8-week-old *huTNFtg* mice, and that this increase is also present in SFs from RA patients^23^. To determine the expression profile of both miR-221 and miR-222 during disease progression, we analyzed their expression in *huTNFtg* SFs obtained at early (3 weeks), intermediate (8 weeks), as well as late (11 weeks) disease stages, and compared them to SFs from WT control mice. Both miR-221 and miR-222 were shown to be upregulated from the early stage of disease and this up-regulation was further augmented with progression of disease (Fig.1A and B).

**Figure 1:**
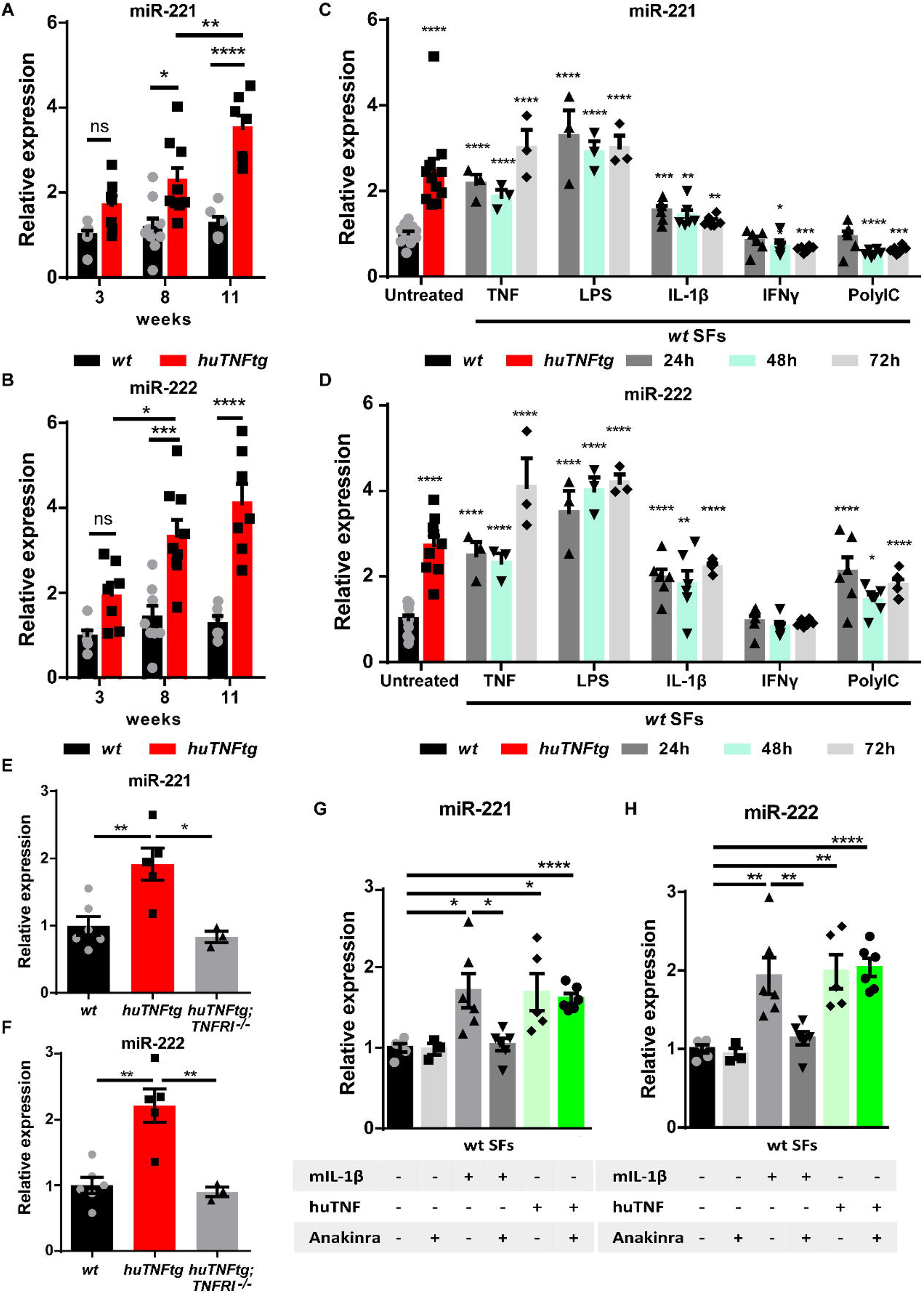
Regulation of miR-221/222 levels in SFs under arthritogenic signals. A-B) Expression levels of miR-221 and miR-222 from cultured SFs, as determined by qRT-PCR at different time points of disease of *huTNFtg* mice, as well as control wild-type mice (WT). Expression levels were normalized to the levels seen in 3 week-old WT mice (n = 6-9). C-D) Quantification of miR-221 and miR-222 levels in cultured WT SFs after 24h, 48h and 72h stimulation with TNF, LPS, IL1-β, IFN-γ, PolyIC (n = 3-13). WT unstimulated SFs served as reference. E-F) miR-221 and -222 levels in cultured SFs from *huTNFtg* and *huTNFtg*;*TNFRI*^-/-^ mice (n = 3-6). WT SFs served as reference for normalization. G-H) miR-221 and -222 expression analysis in cultured WT SFs after TNF or IL-1β stimulation in the absence or presence of anakinra (n = 3-6). Expression was normalized to the levels detected in WT unstimulated SFs. In all expression analysis experiments for miR-221 and -222, *u6* was used as a housekeeping gene. Data represent mean ± SEM. *p < 0.05, **p < 0.01, ***p < 0.001, ****p < 0,0001, ns = not significant.

Previous studies have shown that miR-221 and miR-222 expression is induced by LPS and TNF in macrophages and that they are implicated in LPS tolerance and endotoxemia^28,34,35^. To understand the tissue- and disease-specific regulation of miR-221 and miR-222 in synovial fibroblasts, we stimulated SFs with an array of inflammatory stimuli, such as TNF, LPS, IL-1β, IFN-γ and PolyIC, and analyzed microRNA expression 24, 48 and 72 hours post-induction (Fig. 1C and D). To validate the response of SFs to the inflammatory signals we measured the expression of known downstream targets (Fig. S1A and B). As shown in Fig. 1C and D, TNF, LPS and IL-1β led to consistent upregulation of both miR-221 and miR-222 in SFs at all time points examined. To further characterize the requirement of upstream TNF-signaling for the induction of miR-221 and -222 in arthritogenic fibroblasts, SFs were isolated from *huTNFtg* mice lacking TNFR1, and miR-221/222 levels were measured. These mice do not develop any RA pathology and the two microRNAs were found not to be induced in the absence of TNFR1 (Fig. 1E and F). Additionally, we analyzed whether the TNF-mediated induction of miR-221/222 depends on IL-1β signaling. Since IL-1β signaling is reported to act both independently or downstream of TNF in RA and SFs^36–41^, we treated WT SFs with anakinra (an IL-1 receptor antagonist) and then stimulated them with TNF. Anakinra successfully inhibited downstream IL-1β signaling (Fig. S1C) but did not abrogate miR-221/222 induction by TNF (Fig. 1G-H). Thus, TNF may upregulate miR-221/222 independently of IL-1β.

### Mesenchymal cell overexpression of miR-221 and miR-222 exacerbates arthritis in *huTNFtg* mice

In order to study the *in vivo* role of miR-221/222 in SFs, we generated a transgenic mouse model, *TgColVI-miR-221/222*, overexpressing these two microRNAs under the ColVI promoter (Fig.2A), that is known to target cells of mesenchymal origin in the joints, including the SFs^8^. Overexpression of miR-221 and miR-222 was verified through qRT-PCR analysis in various tissues (Fig. S2A and B). Fibroblast-specific overexpression of miR-221 and miR-222 was verified in *ex vivo* cell cultures of SFs, intestinal mesenchymal cells (IMCs) and lung fibroblasts (LFs) (Fig. S2C and D), whereas no overexpression could be observed in cells of hematopoietic or epithelial lineage (Fig. S2E and F).

To explore the role of the two microRNAs in the pathogenesis of arthritis, we crossed the *TgColVI-miR-221/222* with the *huTNFtg* mice and monitored disease progression. Histopathological analysis at 8 weeks of age revealed exacerbated arthritis, as observed by enhanced synovial hyperplasia (pannus formation), cartilage destruction and number of osteoclasts in the double *huTNFtg;TgColVI-miR-221/222* transgenic mice in comparison to the *huTNFtg* controls (Fig.2B). Moreover, bone morphometric analysis using microCT revealed greater bone erosions as indicated by decreased bone volume, decreased trabecular thickness and increased trabecular separation (Fig.2C-F). Interestingly, comparison of *TgColVI-miR-221/222* to control mice (WT) revealed significantly decreased trabecular thickness, as well as a tendency for decreased bone volume and increased trabecular separation (Fig. 2D-F, Fig. S3), suggesting a potential a priori function of the two microRNAs in bone physiology. miR-221/222 overexpression in the joints of *huTNFtg;TgColVI-miR-221/222* mice did not lead to significant increase of inflammatory infiltrations, apart from an increase in CD8^+^ T cells (Fig.3A, Fig. S4A and B).

**Figure 2:**
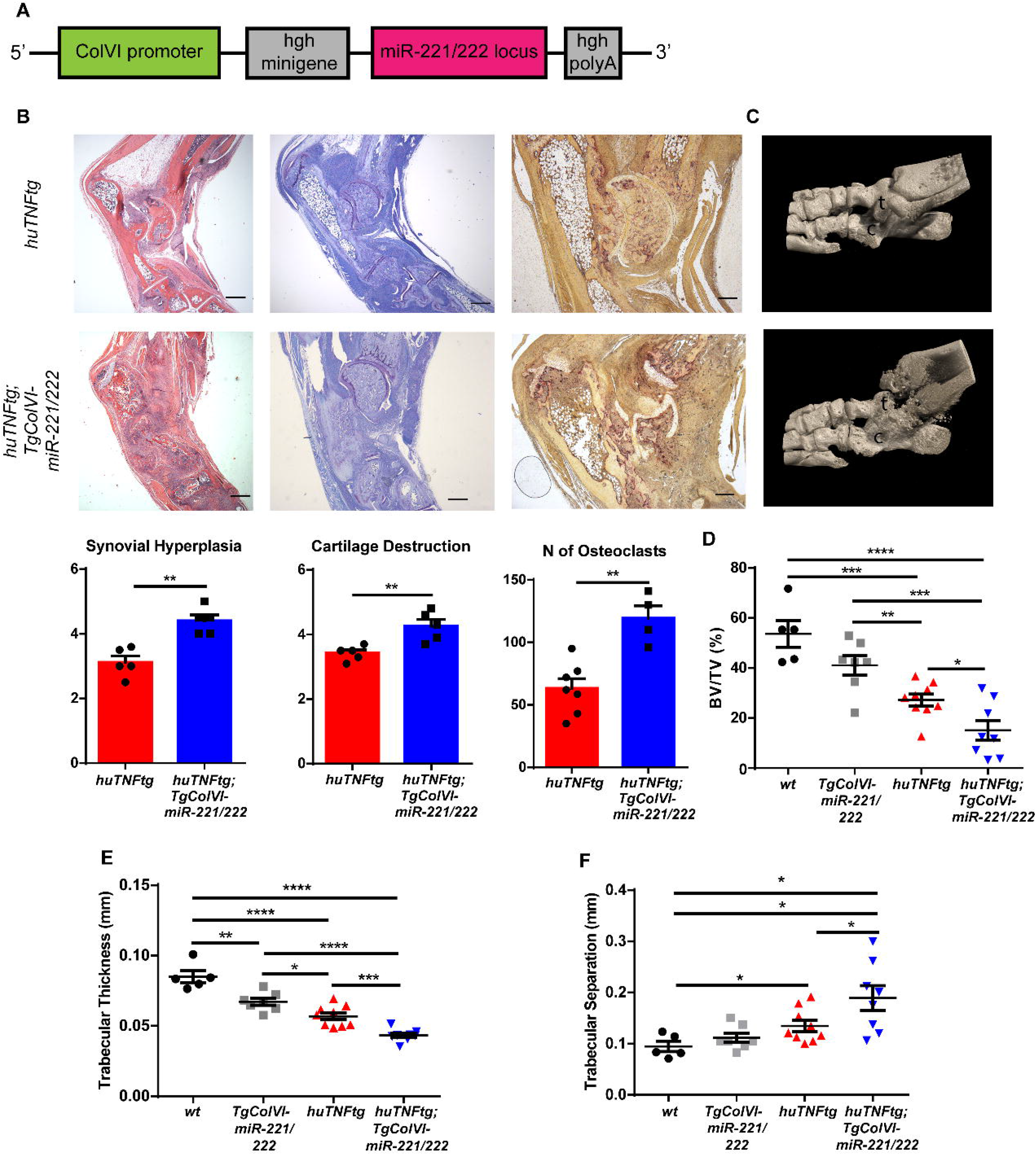
Mesenchymal miR-221/222 overexpression leads to worse arthritis manifestations in *huTNFtg* mice. A) miR-221/222 were cloned under the ColVI promoter to target expression in cells of mesenchymal origin. B) Representative histological images of H&E, TB and TRAP stained ankle joint sections and histological score of synovial hyperplasia, cartilage destruction and osteoclast numbers of 8 week-old *huTNFtg* (n=5-7) and *huTNFtg;TgColVI-miR-221/222* mice (n = 4-5). t: talus, c: calcaneous. C) Representative microCT images of the ankle joint area of 8 week-old *huTNFtg* and *huTNFtg;TgColVI-miR-221/222* mice. D-F) Quantification of bone erosions measuring the following parameters: decreased bone volume, decreased trabecular thickness and increased trabecular separation by microCT analysis in the ankle joints of WT (n = 5), *TgColVI-miR-221/222* (n = 7), *huTNFtg* (n = 9) and *huTNFtg;TgColVI-miR-221/222* mice (n = 8). Scale bars: 600μm and 300μm. Data represent mean ± SEM. *p < 0.05, **p < 0.01, ***p < 0.001, ****p < 0,0001, ns = not significant.

**Figure 3:**
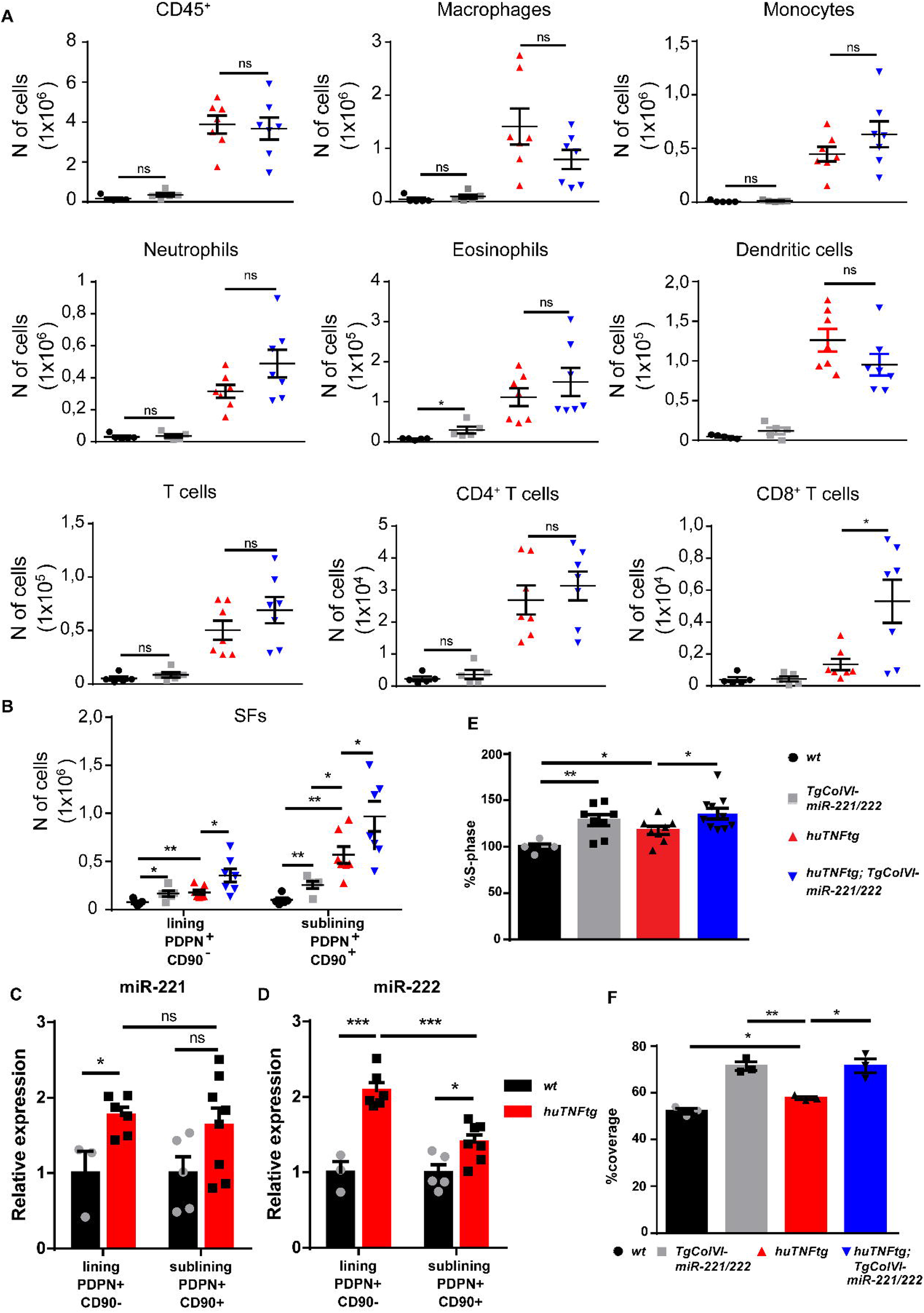
miR-221/222 overexpression leads to fibroblast expansion. A) Infiltration of CD45^+^ cells, macrophages, monocytes, neutrophils, eosinophils, dendritic cells, CD4^+^ T cells and CD8^+^ T cells in the ankle joints of 8 week-old WT (n = 5), *TgColVI-miR-221/222* (n = 5), *huTNFtg* (n = 7) and *huTNFtg;TgColVI-miR-221/222* mice (n = 7) quantified by FACS analysis (from two independent experiments). B) Lining PDPN^+^ CD90^+^ and sublining PDPN^+^CD90^+^ SF number quantification in the ankle joints of 8 week-old WT (n = 5), TgColVI-miR-221/222 (n = 5), *huTNFtg* (n = 7) and *huTNFtg;TgColVI-miR-221/222* mice (n = 7) by FACS analysis (from two independent experiments). C-D) miR-221 and -222 levels in freshly sorted lining PDPN^+^CD90^-^ and sublining PDPN^+^CD90^+^ SFs in the ankle joints of 8 week-old WT and *huTNFtg* mice (n = 3-8). E) % fraction of cultured SFs that are in S-phase of the cell cycle from 8 week-old WT (n = 5), *TgColVI-miR-221/222* (n = 8), *huTNFtg* (n = 8) and *huTNFtg;TgColVI-miR-221/222* mice (n = 10) quantified by FACS analysis (from three independent experiments). F) % area that was covered by cultured fibroblasts 24h after a wound was performed (n = 3). A representative experiment out of three is presented. All comparisons were performed using WT SFs as a reference sample. Data represent mean ± SEM. *p < 0.05, **p < 0.01, ***p < 0.001, ****p < 0,0001, ns = not significant.

To examine the effect of miR-221/222 overexpression in the physiology of synovial fibroblasts, we focused on the analysis of known fibroblast subpopulations in the joints of the *huTNFtg;TgColVI-miR-221/222* mice. The synovium of the joints consists of two compartments, the lining and the sublining, that are characterized by the expression of specific markers. Recent elegant studies in human RA attributed differential functions to these two layers with PDPN^+^CD90^-^ fibroblasts of the lining layer (LLSFs) being destructive, whereas the PDPN^+^CD90^+^ cells of the sublining layer (SLSFs) being proinflammatory^42^. Similar to human RA, both LLSFs and SLSFs were expanded in *huTNFtg* mice compared to WT, as indicated by FACS analysis (Fig.3B, Fig. S4C). Notably, miR-221/222 overexpression in arthritis rendered both layers more hyperplastic in *huTNFtg;TgColVI-miR-221/222* mice compared to *huTNFtg* mice. It is worth mentioning that miR-221/222 overexpression alone was sufficient to increase the number of fibroblasts of the LL and SL in *TgColVI-miR-221/222*. (Fig.3B). To define which SF subpopulation responds to inflammatory signals produced by the arthritogenic microenvironment and upregulates miR-221 and miR-222 expression, we sorted LL and SL fibroblasts and measured miR-221 and -222 expression. As depicted in Fig. 3C and D, an increase of miR-221 and -222 levels is observed, mainly, in fibroblasts of the LL and to a lesser extent in those of the SL.

Finally, primary SFs isolated from the joints of *huTNFtg;TgColVI-miR-221/222* mice were functionally characterized *ex vivo*. In agreement with the *in vivo* data, both WT and *huTNFtg* SFs overexpressing miR-221/222 exhibited increased proliferation and wound healing potential compared to their respective controls (Fig.3E and F). Collectively, these results establish a pro-proliferative function of miR-221/222 on the synovial fibroblasts.

### Overexpression of miR-221/222 in SFs results in activated pathways involved in cell cycle and repressed pathways linked to ECM remodeling

To identify downstream targets of miR-221/222 and unveil the molecular events that mediate the effect of their overexpression, we performed RNA sequencing and analyzed the expression profile of SFs from WT, *TgColVI-miR-221/222, huTNFtg* and *huTNFtg;TgColVI-miR-221/222* mice. The different genotypes are presented with distinct expression signatures and grouped according to genotype after performing Principal Component Analysis (PCA) (Fig. S5A). We found a significant number of genes deregulated in all conditions compared to WT SFs, as well as an increase in deregulated genes due to miR-221/222 overexpression in the arthritogenic environment of *huTNFtg* mice (Fig. S5B and C). To assess the effect of miR-221/222, we focused on genes upregulated (n=399) and downregulated (n=449) in *huTNFtg;TgColVI-miR-221/222* SFs compared to *huTNFtg* (Fig. 4A). KEGG pathway analysis of significantly differentially expressed genes showed strong over-representation of pathways involved in DNA replication and cell cycle progression in the group of upregulated genes, while functional terms such as ECM function and AKT signaling were enriched among the downregulated ones (Fig.4A).

**Figure 4:**
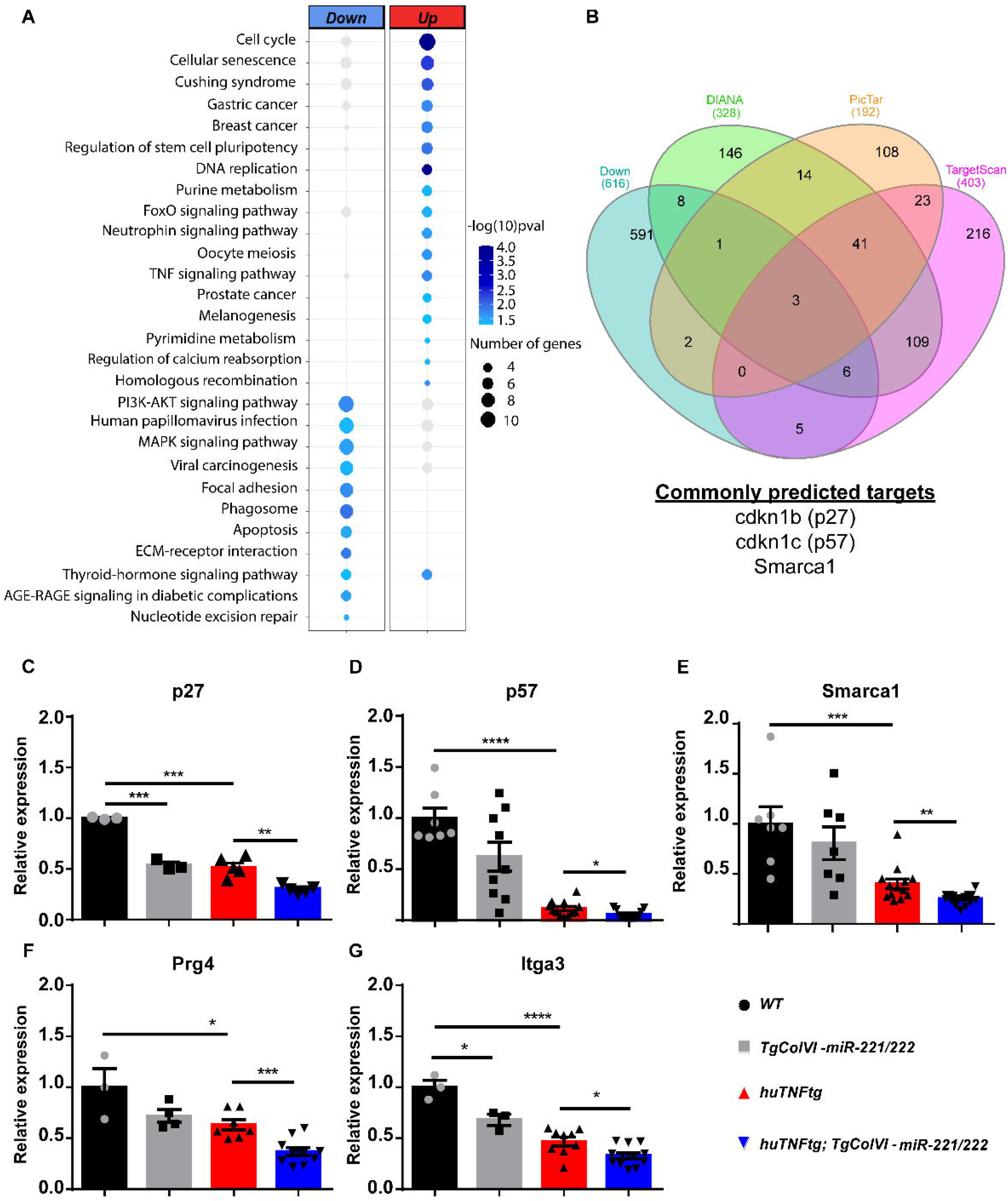
miR-221/222 regulate cell cycle signaling and ECM-related pathways in arthritis. A) Bubble plot of enriched KEGG pathways in the up (red) and down-regulated genes (blue) originated from the comparison of *huTNFtg;TgColVI-miR-221/222* and *huTNFtg* bulk RNA-seq profiles. The color of the bubble signifies the statistical significance, while the size denotes the number of up/down regulated genes found in the enriched term. B) Venn diagram showing the overlap of genes predicted as miR221/222 targets from DIANA, PicTar and TargetScan and downregulated genes from *huTNFtg* compared to WT and *huTNFtg;TgColVI-miR-221/222* compared to *huTNFtg* SFs. C-G) Expression analysis as defined by qRT-PCR of p27, p57, Smarca1, Prg4 and Itga3 in cultured SFs from 8 week-old WT, *TgColVI-miR-221/222, huTNFtg* and *huTNFtg;TgColVI-miR-221/222* mice (n = 3-16). WT SFs were used as a reference. Data from 2-4 independent experiments. In all experiments B2m was used as housekeeping gene for normalization. Data represent mean ± SEM. *p < 0.05, **p < 0.01, ***p < 0.001, ****p < 0,0001, ns = not significant.

To uncover direct targets of miR-221/222, we cross-referenced downregulated genes in SFs from *huTNFtg* compared to WT and from *huTNFtg;TgColVI-miR-221/222* compared to *huTNFtg* mice with their predicted targets using three different tools (DIANA-microT-CDS, Targetscan and Pictar) (Fig.4B). Among these potential target candidates, we found p27, p57 and Smarca1. p27 and p57 are cell cycle inhibitors and have been previously validated as miR-221/222 targets^24,26^, while Smarca1 (a component of the NURF complex involved in chromatin remodeling^43^ is a new predicted target. To verify their regulation by miR-221/222, the expression levels of all targets were determined in SFs isolated from *huTNFtg;TgColVI-miR-221/222* mice, as well as control mice, using qRT-PCR. All p27, p57 and Smarca1 were found to be downregulated in SFs from *huTNFtg;TgColVI-miR-221/222* mice compared to *huTNFtg* mice, as well as in SFs from *huTNFtg* mice compared to SFs from WT mice (Fig. 4C-E). Together our data, from RNA sequencing and subsequent analysis, suggest that miR-222/221 target specific cell cycle inhibitors (p27 and p57), in order to induce proliferation of fibroblasts. Apart from these two targets that play a role in cell proliferation, we also identified Smarca1, a novel predicted miR-221/222 target involved in chromatin dynamics. ECM related genes such as Prg4 and Itga3 were found to be downregulated in *huTNFtg* SFs (Fig. 4F and G). Interestingly, miR-221/222 overexpression resulted in enhanced downregulation of these genes, probably through indirect mechanisms as they were not predicted to be direct targets.

We further analyzed if IL-6 expression levels in SFs were regulated by miR-221/222 overexpression, as previously reported^31^. We did not observe any changes neither in the IL-6 protein levels in supernatants of naïve or TNF stimulated SFs nor in IL-6 RNA levels in SFs from miR-221/222 overexpressing mice (Fig. S6A and B).

Additionally, TNF has been reported to be a direct target of miR-221 and miR-222 in sepsis in macrophages^28,34^. However, in the *huTNFtg* model of arthritis, this could not apply as the endogenous 3’UTR of the *huTNF* gene is replaced by the 3’UTR of the human *β-globin* gene^3^. Nevertheless, we checked if there are indirect signals that stem from miR-221/222 overexpression in SFs and affect the already deregulated expression of *huTNF*. RNA levels of *huTNF* were not altered significantly in SFs from *huTNFtg;TgColVI-miR-221/222* mice compared to *huTNFtg* (Fig. S6C). Moreover, no mTNF protein was detected in the sera or supernatants of cultured SFs from *huTNFtg* and *huTNFtg;TgColVI-miR-221/222* mice. Furthermore, in inflamed joints, SFs become activated and act as recruiters of leukocytes by increasing the expression of adhesion molecules. To examine whether miR-221/222 overexpression rendered fibroblasts more activated, we measured the expression of activation markers, such as ICAM-1 and VCAM-1, but no differences were observed due to miR-221/222 overexpression (Fig. S6D and E). It is therefore evident that miR-221/222 overexpression shifts the gene expression of arthritogenic SFs to a more proliferative and less ECM-producing signature, without any evidence of switching to a more pro-inflammatory or activated state.

### Total deletion of miR-221/222 ameliorates arthritis in *huTNFtg* mice

To assess the therapeutic potential of miR-221/222 downregulation, we studied the impact of miR-221 and miR-222 deletion in arthritis. Crossing mice carrying targeted allele of miR-221/222 with loxP sites (*miR-221/222 f/f*) with a *Deleter*-*Cre*^44^ mouse line led to the generation of complete knockout mice, referred to as *miR-221/222* ^-/-^. Deletion of miR-221/222 in SFs of *miR-221/222* ^-/-^ mice was confirmed by qRT-PCR (Fig. S7A and B). *miR-221/222* ^-/-^ mice developed normally, did not exhibit any obvious phenotypic defects and were crossed with *huTNFtg* mice to ablate miR-221/222 during arthritis. *huTNFtg;miR-221/222* ^-/-^ mice at 9 weeks of age presented with decreased synovial hyperplasia, cartilage destruction and number of osteoclasts, as seen in histological analysis compared to *huTNFtg* (Fig.5A). Additionally, miR-221/222 deletion in arthritis did not alter the inflammatory influx (Fig.S8), but rather led to inhibition of fibroblast expansion of LL and SL in the joints (Fig.5B). Ex vivo, functional characterization of arthritic SFs in the absence of miR-221/222 showed decreased proliferative capacity than SFs from *huTNFtg* mice (Fig.5C), confirming the control of cell proliferation by these two microRNAs. p27 and Smarca1 levels (and not p57) were partially restored (Fig. 5D-F), verifying their regulation by miR-221/222, although alternative or feedback mechanisms might also exist preventing the complete restoration of the expression levels of these genes and complete disease protection of the *huTNFtg;miR-221/222* ^-/-^ mice.

**Figure 5:**
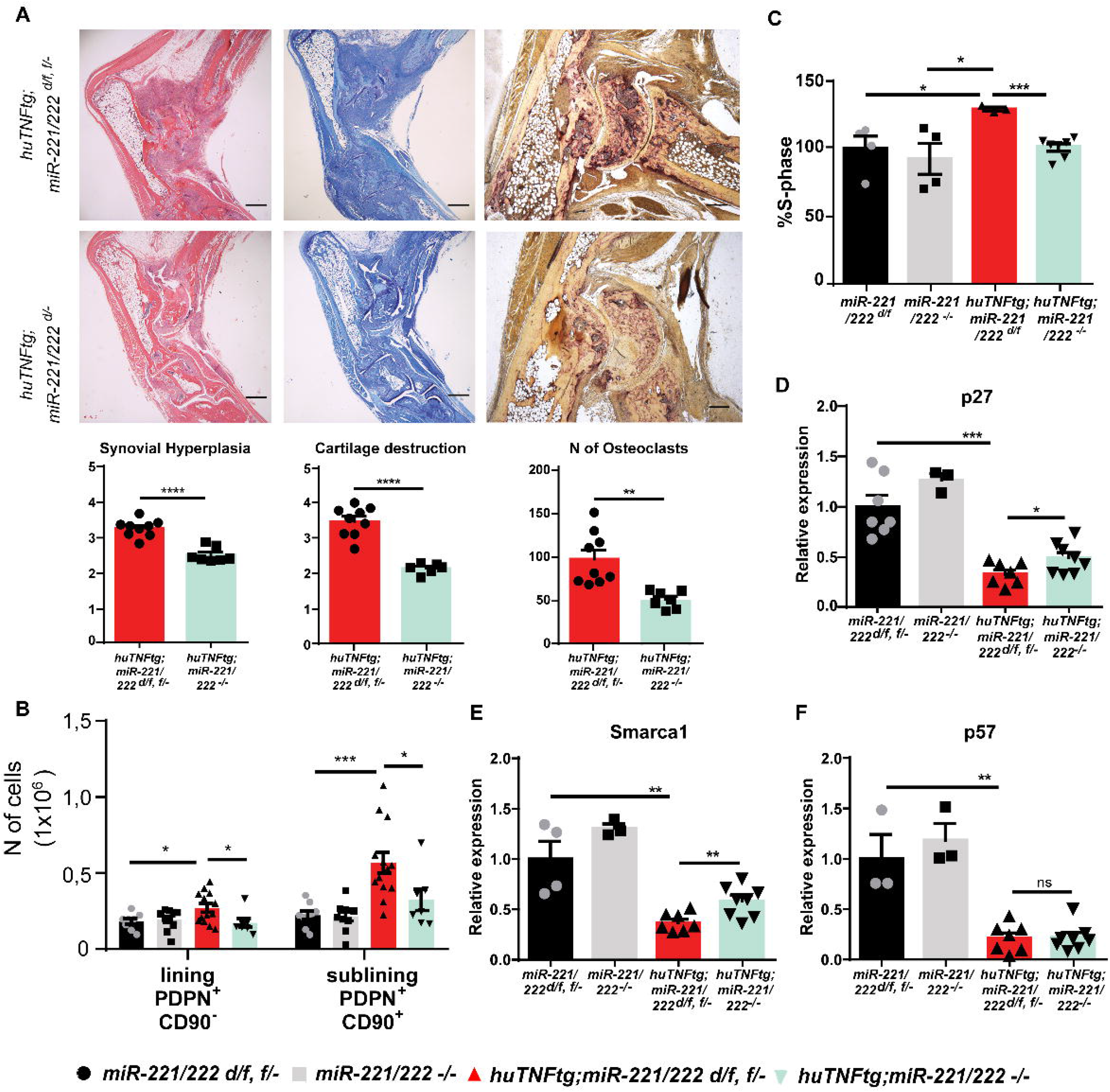
Deletion of miR-221/222 ameliorates arthritis in *huTNFtg* mice. A) Representative histological images of H&E, TB and TRAP stained ankle joint sections and histological score of synovial hyperplasia, cartilage destruction and osteoclast numbers from 9 week-old *huTNFtg;miR-221/222 d/f*, f/- (n= 9) and *huTNFtg*;*miR-221/222*^*-/-*^ mice (n = 6-7). B) Lining PDPN^*+*^CD90^-^ and sublining PDPN^*+*^CD90^*+*^ SF number quantification in the ankle joints of 9 week-old *miR-221/222 f/-, d/f* (n = 8-9), *miR-221/222* -/- (n = 10), *huTNFtg;miR-221/222 f/-, d/f* (n = 13) *huTNFtg*;*miR-221/222* - /- mice (n = 7) by FACS analysis (from three independent experiments). C) % fraction of cultured SFs that are in S-phase of the cell cycle from 9 week-old *miR-221/222 d/f* (n = 4), *miR-221/222* -/- (n = 4), *huTNFtg;miR-221/222 d/f* (n = 3) and *huTNFtg;miR-221/222 -/-* (n = 7) (from two independent experiments). *miR-221/222 d/f* SFs was used for normalization. D-F) Expression analysis as defined by qRT-PCR of p27, p57 and Smarca1 in cultured SFs from 9 week-old *miR-221/222 f*/-, *d/f, miR-221/222 -*/-, *huTNFtg*;*miR-221/222 d/f* and *huTNFtg*;*miR-221/222* -/- (n = 3-8, from two independent experiments). Expression of *miR-221/222 d/f, f*/- SFs was used for normalization. In all experiments B2m was used as housekeeping gene for normalization. Scale bars: 600μm and 300μm. Data represent mean ± SEM. *p < 0.05, **p < 0.01, ***p < 0.001, ****p < 0,0001, ns = not significant.

### miR-221/222 gene activity marks the pathogenic clusters of expanding intermediate and lining compartment of SFs in arthritis

Our next step was to define miR-221/222 gene activity scores at a single cell level in arthritogenic SFs. Chromatin accessibility analysis of previously generated scATAC-seq data from ankle joint fibroblasts of *WT* and *huTNFtg* mice^45^ revealed the emergence and expansion of a pathogenic intermediate cluster and a destructive one in the lining (Fig. 6A, Fig. S9A), along with clusters found physiologically in *WT* mice. As can be seen, miR-221/222 locus is more accessible in the pathogenic intermediate and destructive lining clusters (Fig. 6B, Fig. S9B-C), marking the aberrant expansion of these clusters. To gain insight into which regulatory elements may control miR-221/222 expression in arthritogenic SFs, correlation analysis between chromatin accessibility and gene activity scores was performed in the regulatory space of miR-221/222. Inferred linkages between accessible chromatin (peaks) and miR-221/222 genetic locus, revealed transcription factor binding sites (TFBSs) of previously reported positive regulators such as Nrf2 (Nfe2l2), AP-1, NF-κB and Bach1^45^, to be enriched in the regulatory regions associated with miR-221/222 locus (Fig. 6C, Fig. S9D). Overall, the increased accessibility of miR-221/222 locus in the activated and expanding intermediate cluster, as well as in the destructive cluster of the lining layer in arthritis corroborates the role of these two microRNAs in promoting proliferation and expansion of activated clusters of SFs in disease.

**Figure 6:**
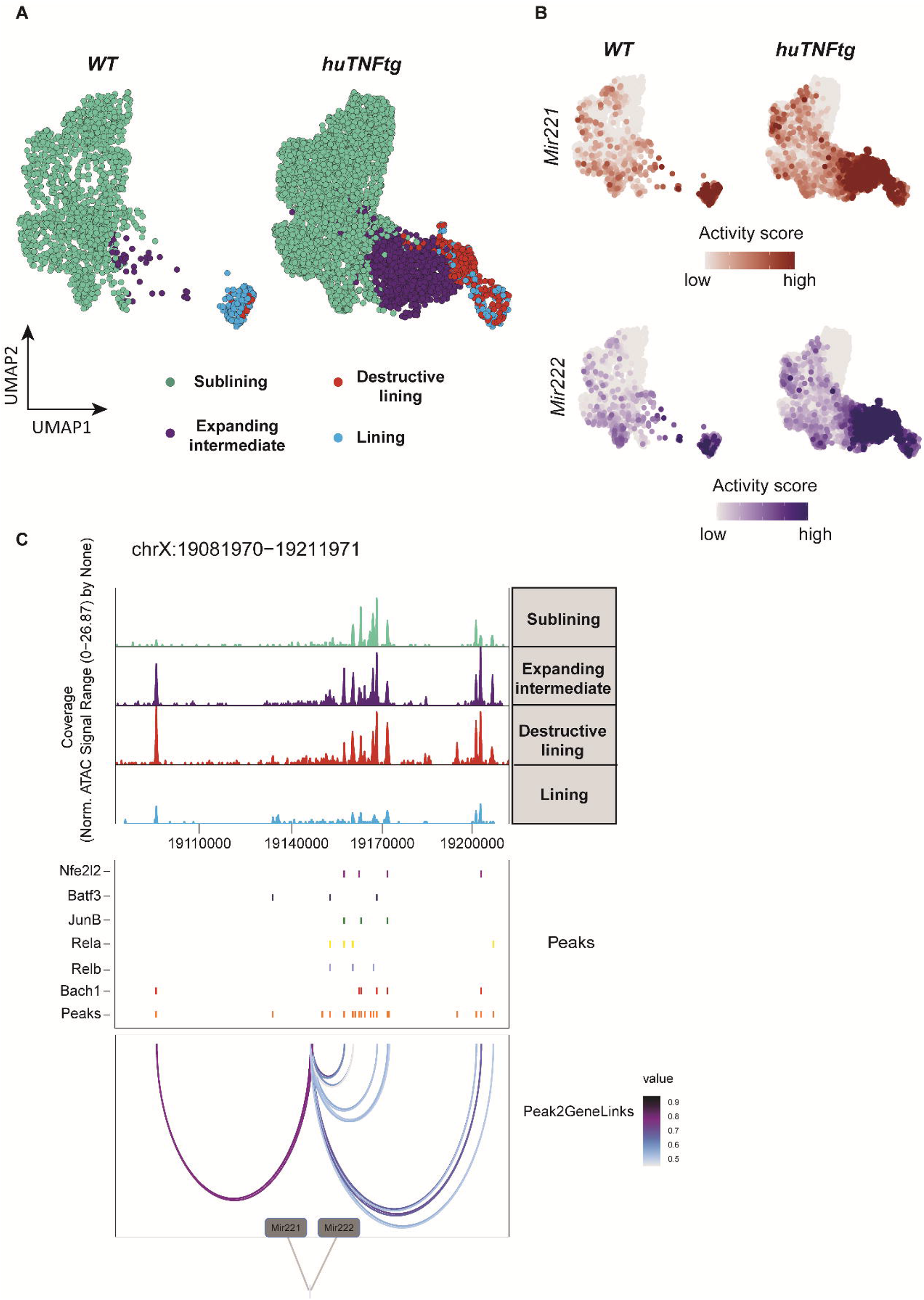
miR-221/222 gene activity is increased in the pathogenic subclusters of the expanding intermediate and lining compartment. A) UMAP projection of 6,046 synovial fibroblast nuclei obtained by scATAC-seq from *WT* and *huTNFtg* samples. Cells are grouped in four categories: Sublining (light green), Expanding intermediate (purple), Lining (light blue) and Destructive lining (red). B) Feature plots, in UMAP space, depicting gene activity scores of Mir221 (red) and Mir222 (blue) in *WT* and *huTNFtg* samples. C) Genome accessibility track visualization of the extended regulatory space of Mir221 and Mir222 (chrX:19,081,970-19,211,971), with TF binding site and peak-to-gene linkage information, in *huTNFtg* samples. Upper, the genome track shows increased accessibility in expanding intermediate and destructive lining clusters. Middle, all reproducible peaks are shown, coupled with annotated CISBP binding information for Bach1, Rela, Relb, Nfe2l2 (Nrf2), Batf3 and JunB TFs. Lower, putative regulatory linkages between Mir221-Mir222 genes and reproducible peaks are illustrated. Links between genes and peaks are colored by correlation (Pearson coefficient) of peak accessibility and gene activity scores.

## Discussion

microRNAs are important fine tuners of gene expression and often exhibit a specific tissue or developmental expression pattern^46^. Deregulation in microRNA expression levels has been linked with diseases such as cancer, viral infection and inflammation^47,48^.

In RA, alteration in expression of several miRNAs has been reported in different cell types in the synovium or in circulating PBMCs (peripheral blood mononuclear cells). miR-146a and miR-155 are among the most highly studied microRNAs and both are upregulated in response to inflammatory signals and implicated in immune responses^18,19,49–51^. More specifically, miR-146a was found upregulated in various RA patient cell types and *in vivo* exogenous double-stranded miR-146a administration in autoantigen driven arthritis in mice led to a reduction in bone erosions through inhibition of osteoclastogenesis without affecting synovial inflammation^52^. Moreover, miR-146a total ablation in *huTNFtg* mice led to increased joint pathology and activation of fibroblasts via TRAF6 downregulation^53^. Although, ex vivo studies had also indicated a protective role for miR-155 by downregulating MMP-1 and MMP-3 expression^54^, miR-155 deficient mice were actually protected against antigen-driven arthritis via the blockade of antigen-specific Th17 polarization. Additionally, miR-155 deficient mice showed less immune-mediated bone erosions via inhibition of osteoclastogenesis in the serum-transfer model of arthritis^55,56^. Despite a study reporting an anti-inflammatory role for miR-23b when overexpressed in radio-resistant cells in CIA and EAE^57^, in the vast majority of the studies, no tissue-specificity for the function of miRNAs in arthritis was considered.

In this study, we try to shed light on the *in vivo* role of the miR-221/222 family of miRNAs in arthritis. In the past, IL-1β had been reported to mediate downstream signaling of TNF/TNFR1 axis^36,37,40,41^. In the present study, we provide evidence that in the arthritic microenvironment of the joints, TNFR1 signaling downstream of TNF and independently of IL-1β can induce miR-221/222 levels in SFs apart from LPS and could suggest that increased TNF and IL-1β levels can serve as signals that lead to miR-221 and miR-222 upregulation specifically in SFs of RA patients. We observed miR-221 and miR-222 induction to reach peak levels in late disease stages and this agrees with a previous study that correlates their levels with disease activity in RA patients^30^. Thus, these microRNAs could serve as biomarkers to predict disease progression in humans.

In RA, proliferation of the fibroblasts consists one of the hallmarks of the aggressive behavior of the pathogenic mesenchyme. This aberrant propagation of the joint synovium is mediated by survival and anti-apoptotic signals in SFs^58^. Although the signals and epigenetic alterations that lead to this aberrant proliferation of SF subsets in RA, even when cultured *ex vivo*, remain under investigation, in this study we uncovered a novel role for miR-221/222 in the mesenchyma, where increased levels of miR-221/222 rendered fibroblasts more proliferative and migratory, even in the absence of an inflammatory trigger. Moreover, a causal role for miR-221/222 involvement in pannus hyperplasia was shown by the total ablation of these two microRNAs in the arthritic mice, that led to decreased disease manifestations due to attenuated proliferation of SFs and decreased numbers of fibroblasts comprising the lining and sublining layers. Also, considering that mesenchymal over-expression of miR-221/222 at naïve conditions (*WT* background) led to a tendency towards worse bone integrity and increased fibroblast populations, suggests that miR-221/222 upregulation and control of target gene expression could have dramatic effects in a specific milieu under local unresolved inflammation.

Analysis of the transcriptional profile of SFs from the *huTNFtg;TgColVI::miR-221/222* mice compared to the *huTNFtg* mice uncovered deregulation in pathways linked to cell cycle and ECM regulation. Cell cycle inhibitors p27 and p57, known miR-221/222 targets, were both identified in our data and were verified by *ex vivo* experiments in SFs. Interestingly, deregulation in ECM components was observed in the expression analysis data due to miR-221/222 overexpression. ECM plays a crucial role in the joints and it does not exist merely to provide support, but also promotes cell communication and motility. Synovial fluid containing hyaluronic acid and lubricin (Prg4) protects from friction and lubricates the cavity area^59^. Interestingly, during miR-221/222 overexpression in arthritis, the expression of ECM-related molecules in SFs was found altered and, more specifically, lubricin and the integrin component Itga3 were downregulated. Lubricin and Itga3 were not predicted as direct targets of miR-221/222, so indirect mechanisms probably mediate this effect. Itga3 expression has been found to be downregulated in PBMCs from OA (osteoarthritis) patients^60^ and a previous study has reported that lubricin absence from the synovium leads to OA and aberrant fibroblast expansion and proliferation^61^. Moreover, a cardioprotective and antifibrotic role of miR-221/222 is observed in cardiac fibroblasts by the downregulation of different types of collagens^62^. So, miR-221/222 could contribute to the catabolic profile of SFs in arthritis by affecting expression of ECM components. Furthermore, the mesenchymal overexpression of miR-221/222 did not render SFs more inflammatory or activated per se as it was supported *by ex vivo* studies in the past^31^. In contrast, our *in vivo* data suggest that these two microRNAs regulate pro-proliferative signals leading to aberrant expansion of pathogenic SFs that in the end renders the arthritic milieu more inflammatory.

Additionally, Smarca1 was identified as a new target of miR-221/222 based on the transcriptional profile of SFs and bioinformatic analysis of predicted targets. Smarca1 participates in the NURF chromatin remodeling complex, that can limit or promote chromatin accessibility, and thus block or activate gene expression correspondingly^43^. NURF complex has been shown to promote cell proliferation, but a role for this complex in RA is missing^63,64^. Recently, a study reported a role for miR-222 in targeting Brg1, a SWI/SNF chromatin remodeling component, leading to repression of inflammatory cytokine expression in sepsis^28^.

Finally, analysis of chromatin accessibility data at a single cell level revealed increased chromatin accessibility in miR-221/222 locus in the activated and expanding intermediate cluster and in the destructive cluster of the lining. This analysis coupled with transcription factor binding site linkage revealed known (such as AP-1 and NF-Kb^27,32^) and novel transcription factors (such as Nrf2 (Nfe2l2) and Bach1), that may play a role in positively regulating the activity of these two microRNAs in arthritis, leading to the expansion of pathogenic and destructive clusters of the intermediate and lining compartments, respectively. Notably, there is a recent study uncovering a role for Nrf2 (Nfe2l2) and Bach1 in promoting NSCLC tumor metastasis^65^. Future experiments may thus elucidate the exact role of these transcription factors in the aggressive, autonomous and migratory character of RA SFs.

The fact that miR-221 and miR-222 are upregulated in SFs from RA patients and that their overexpression is sufficient to worsen the severity of the disease, shows that these miRNAs could be used as potential disease progression biomarkers. Additionally, therapeutic targeting of miR-221/222 could be beneficial, as total deletion of these two microRNAs protected to a certain degree from arthritis progression and the expression of their confirmed targets was partially restored. This partial protection underlines the fact that most probably there are also other mechanisms assuring the suppression of these targets and should be targeted. It is known that miRNA function is often redundant, this means that if a miRNA is missing, other miRNAs can regulate its targets. Further studies are therefore needed, in order to identify additional miRNAs, that target the same genes and assess the potential of their combined downregulation in the development of new treatments. Nonetheless, it is very interesting that during arthritis there are redundant and complementary pathways ensuring expansion of the pathogenic stroma subsets. It is of great interest to dissect which pro-survival and pro-proliferative pathways lead to expansion of pathogenic fibroblast subpopulations, that could constitute novel targets for subpopulation-specific therapeutic intervention.

Collectively, we report that miR-221 and miR-222 lead to enhanced disease progression in a mouse model of inflammatory polyarthritis, due to increased pathogenic synovial fibroblast proliferation under inflammatory conditions, mainly through downregulation of cell cycle inhibitors. Moreover, these microRNAs probably control the expression of chromatin remodeling components such as Smarca1 in SFs in arthritis. Future studies will reveal a specific role for Smarca1 in shaping gene expression in arthritis. Finally, miR-221/222 gene activity is increased in the pathogenic and expanding intermediate and destructive lining clusters. Thus, mesenchymal targeting of miR-221/222 could potentially serve as a therapeutic tool for treating rheumatoid arthritis and their expression could be used as biomarkers for predicting disease outcomes.

## Methods

### Mice

Human TNF transgenic (*huTNFtg*), *TNFR1*^-/-66^ and *Deleter-Cre*^44^ mice have been previously described. *miR-221/222 f/f* were purchased from European Mouse Mutant Archive (EMMA) provided by Helmholtz Zentrum Muenchen (EMMA ID: EM 05507). *Deleter-Cre* were used to generate *miR-221/222*^-/-^ mice. *TgColVI-miR-221/222* mice were generated in Kollias Lab, BSRC Al. Fleming Institute. Briefly, around 1kbp containing the miR-221/222 locus was inserted into the BplI site of intron 2 of hgh minigene in a pBluescript SK (+) vector. Then, the ColVI promoter was inserted upstream of the previous fragment in the SalI – BamHI (blunt) site of the pBluescript vector. The ColVI::miR-221/222 transgene (around 11kbp) was excised as a SalI – NotI fragment for pronuclear microinjections and generation of *TgColVI-miR-221/222* mice. Genetic analysis (by Southern Blot) of the transgenic TgColVI-miR-221/222 mice showed that the transgene is inserted in a head-to-tail direction and in approximately 20 copies.

All mice were maintained in a C57BL/6J or CBA;C57BL/6J genetic background. Mice were maintained under specific pathogen-free conditions in conventional, temperature-controlled, air-conditioned animal house facilities of BSRC Al. Fleming with 12h light/12h dark cycle and received food and water ad libitum.

### Histology

Formalin-fixed, EDTA-decalcified, paraffin-embedded mouse joint tissue specimens were sectioned and stained with haematoxylin-eosin (H&E), Toluidine Blue or Tartrate-Resistance Acid Phosphatase (TRAP) Kit (Sigma-Aldrich). H&E and TB were semi-quantitatively blindly evaluated for the following parameters: synovial inflammation/hyperplasia (scale of 0–5) and cartilage erosion (scale of 0–5) based on an adjusted, previously described method (Mould 2003). TRAP staining of joint sections was performed to measure number of osteoclasts using ImageJ software. Images were acquired with a Nikon microscope, equipped with a QImaging digital camera.

### Microcomputed tomography

Microcomputed tomography (microCT) of excised joints was carried out by a SkyScan 1172 CT scanner (Bruker, Aartselaar, Belgium) at BSRC Al.Fleming Institute following the general guidelines used for assessment of bone microarchitecture in rodents using microCT^67^. Briefly, scanning was conducted at 50⍰kV, 100⍰mA using a 0.5-mm aluminum filter, at a resolution of 5⍰mm/pixel. Reconstruction of sections was achieved using the NRECON software (Bruker) with beam hardening correction set to 40%. The analysis was performed on a spherical volume of interest (diameter 0.54⍰mm) within 73 slides of the trabecular region of calcaneus. Morphometric quantification of trabecular bone indices such as trabecular bone volume fraction (BV/TV%), bone surface density (BS/TV%), trabecular number (Tb. N; 1/mm) and trabecular separation (Tb. Sp; mm) were performed using the CT analyzer program (Bruker). Additionally, 3D images of the scanned area of the samples were generated using the CTvox software (Bruker).

### Cell culture

Primary mouse SFs were isolated from the ankle joints of mice^68^ with the indicated genotypes, were depleted from CD45^+^ cells using Biotinylated anti-mouse CD45 antibody (Biolegend) and Dynabeads Biotin Binder (Invitrogen) according to manufacturer’s instructions and cultured for three to four passages for subsequent experiments.

SFs were seeded at a concentration of 8 × 10^5^ on a 60mm plate, starved overnight, stimulated the next day with either huTNF (10ng/ml), LPS (100ng/ml), mIL-1β (10ng/ml), IFN-γ (10ng/ml) or PolyIC (20μg/ml) and collected at the indicated time points.

In experiments using the IL-1 inhibitor (Anakinra), pretreatment of cells with Anakinra at a concentration of 10μg/ml was performed for 4⍰h before stimulation with mIL-1β (10ng/ml) or huTNF (10ng/ml) and subsequent analysis.

### FACS analysis

FACS analysis and sorting of SF subpopulations were performed by removing ankle joints, cutting them into pieces and being digested using 1000U/ml Collagenase IV (Sigma-Aldrich) in DMEM for 60 minutes at 37°C. The cell suspension was centrifuged, resuspended in FACS buffer (PBS with 0.5% FBS, 0.05% sodium azide and 2mM EDTA) and cells were counted. The anti-Fc Receptor (anti-CD16/32) antibody (Biolegend) was used to prevent non-specific binding. For stainings, 1-2 million cells were incubated with the following antibodies: PE-conjugated anti-CD11b (eBioscience), APC/Cy7- or A700-conjugated anti-CD45 (Biolegend), APC-conjugated anti-MHCII, PE/Dazzle594-conjugated anti-CD64, APC/Cy7-conjugated anti-CD24, FITC-conjugated anti-Ly6C, PE/Cy7-conjugated anti-CD11c (Biolegend), PE-conjugated anti-B220 (eBioscience), PE/Cy7-conjugated anti-CD3, A700-conjugated anti-CD4 (eBioscience), APC-conjugated anti-CD8 (Biolegend), A488-conjugated anti-CD90.2, PE-conjugated anti-CD31, PE/Cy7-conjugated anti-PDPN, Biotinylated anti-Ly6G, streptavidin-conjugated PE/Cy5 (Invitrogen), PE-conjugated anti-ICAM-1 (BD Pharmigen) and A647-conjugated anti-VCAM-1 (Biolegend). Zombie Green or NIR (Sigma) and DAPI (Invitrogen) were used for live/dead exclusion. Analysis was performed using a FACS Canto II Flow cytometer (BD Biosciences) and FACS Diva (BD Biosciences) or FlowJo software (FlowJo, LLC) and cell sorting was performed using a FACS Aria III Cell Sorter (BD Biosciences).

### Cell cycle analysis

Cell cycle was detected by flow cytometry. Briefly, 4 × 10^5^ cells were seeded onto 60 mm plates and incubated for 24 hours. The cells were then harvested and washed with PBS. Next, the pellet was resuspended and fixed in 70% prechilled ethanol for 30min at 4°C. The cells were washed again with PBS followed by treatment with RNase for 30min at 37°C and addition of 200 μl staining solution with propidium iodide into the pellet. The final mixture was analyzed by flow cytometry.

### ELISA

Supernatants were collected at 0, 24 and 48h and IL-6 quantification was performed using the mouse IL-6 Duo-Set ELISA kit, according to the manufacturer’s instructions (R&D Systems).

### Wound healing assay

SFs were plated to form a monolayer on wells of a 24-well plate, serum starved overnight and wounded with 200 μl pipette tips the next day. The culture dishes were washed 3 times with 1XPBS to remove detached cells, and the remaining cells were grown in DMEM containing 10% FBS. After 24 hours of incubation, wound healing potential was quantified by counting the surface that cells had migrated, proliferated and covered using ImageJ Software analysis tool.

### RNA Isolation and qRT-PCR

RNA was isolated from SFs, intestinal mesenchymal cells, spleenocytes and epithelial cells using the RNeasy or the miRNeasy mini kit (QIAGEN), according to manufacturer’s instructions. Isolated RNA was subsequently used either for 3⍰RNA-seq sequencing and analysis, or for construction of cDNA, using the MMLV Reverse Transcriptase (Promega) or the Taqman MicroRNA Reverse Transcription kit (Applied Biosystems). The cDNA was subsequently used for qRT-PCR using the Platinum SYBR-Green qPCR SuperMix (Invitrogen) or for detection of microRNA levels the Taqman gene expression mastermix (Applied Biosystems) and the relative Taqman microRNA probes. The CFX96 Touch Real-Time PCR Detection System (Biorad) was used. Quantification was performed with the DDCt method. Primer sequences (5⍰-3⍰) are provided (see Table S1).

### 3⍰ RNA-Seq sequencing

The quantity and quality of RNA samples were analyzed using Agilent RNA 6000 Nano kit with the bioanalyzer from Agilent. RNA samples with RNA Integrity Number (RIN) > 7 were used for library construction using the 3⍰ mRNA-Seq Library Prep Kit Protocol for Ion Torrent (QuantSeq-LEXOGEN™) according to manufacturer’s instructions. DNA High Sensitivity Kit in the bioanalyzer was used to assess the quantity and quality of libraries, according to manufacturer’s instructions (Agilent). Libraries were then pooled and templated using the Ion PI™ IC 200 Kit (ThermoFisher Scientific) on an Ion Proton Chef Instrument or Ion One Touch System. Sequencing was performed using the Ion PI™ Sequencing 200 V3 Kit and Ion Proton PI™ V2 chips (ThermoFisher Scientific) on an Ion ProtonTM System, according to manufacturer’s instructions.

### 3’ RNA-seq bioinformatics analysis

Quality of the FASTQ files (obtained after Ion Proton sequencing) was assessed by FastQC, following the software recommendations. Alignment of sequencing reads to the mouse reference genome was performed by the software HISAT2 (version 2.1.0) with the mm10 version of the reference genome. The raw bam files were summarized to read counts table using FeatureCounts (version 1.6.0) and the gene annotation file mus_musculus.grcm38.92.gtf from Ensembl database. The resulting gene counts table was subjected to differential expression analysis (DEA) utilizing the R package DESeq2. Differentially expressed genes were identified after setting the following thresholds: pvalue < 0.05 and |log2FC| > 1. Functional enrichment analysis of the up and downregulated genes for the contrasts of interest was performed with the web version of the tool enrichR. Enriched KEGG pathways (KEGG Mouse 2019) were selected by setting the following thresholds : pvalue < 0.05 and gene count > 2.

### scATAC-seq bioinformatics analysis

Analysis of scATAC-seq datasets was conducted by using the cellranger (10X genomics) and ArchR suite as previously described^45,69^. Briefly, BCL files were converted to FASTQ files, aligned to UCSC mm10 reference genome, and *WT* and *huTNFtg* samples were aggregated and counted using 500-bp bins (tiles). Latent Semantic Indexing (LSI), graph-based clustering (louvain), and UMAP dimensionality reduction was applied accordingly. Gene activity scores (predictions of the level of expression of each gene) were computed as described in ArchR. Gene scores were scaled to 10,000 counts and log-normalized. To assign scATAC-seq cell type identiy, gene-activity scores and scRNA-seq gene expression^45^ were aligned directly using a two-stage Canonical Correlation Analysis (CCA). Non-fibroblast cells were excluded to result in 6,046 SFs cells, which were reanalyzed as described above. The integration process between scATAC-seq and scRNA-seq SFs labeled the scATAC-seq cells according to 4 SF subpopulations (homeostatic, intermediate, lining, and destructive lining, that were visualized in a UMAP 2D space. Peak calling was applied in two pseudo-bulk replicates across all SFs^69^, and then merged using an iterative overlap peak-merging^70^. To identify enriched motifs in single-cell resolution, chromVar was used^71^. Positive TF regulators were identified by correlating TF motif accessibility with integrated TF gene expression (Pearson Correlation Coefficient > 0.5 and P-adjusted value < 0.05). Finally, to identify regulatory links between accessible regions and active genes, peak to gene linkages were inferred using correlation analysis between enhancer accessibility and gene activity scores^69^.

### Statistical analysis

All experiments were performed at least 3 times. Data are presented as mean⍰±⍰SE. Student’s t-test (parametric, unpaired, two-sided) or two-way ANOVA were used for evaluation of statistical significance using GraphPad 6 software. Statistical significance is presented as follows: * p < 0.05, ** p < 0.01, *** p < 0.001, **** p < 0.0001.

### Study Approval

All experiments were approved by the Institutional Committee of Protocol Evaluation in conjunction with the Veterinary Service Management of the Hellenic Republic Prefecture of Attika according to all current European and national legislation.

## Author’s Contributions

GK and FR designed the study and interpreted the experimented results. FR, CT, DK, DP, AP, MS, AL, PH, TM and YP performed the experiments and data analysis. NK, MD, MA and MF contributed to data interpretation. FR drafted the manuscript and the figures with contribution from CT and DK. GK edited and finalized the manuscript. All authors were involved in critically revising the final manuscript. All authors read and approved the final manuscript.

## Acknowledgments

We thank Dr. Aikaterini Nanou for critically reviewing the manuscript. We also thank Lida Iliopoulou, Anna Katevaini, Spiros Lalos, Panos Athanasakis, Michalis Meletiou and Kleopatra Dagla for excellent technical assistance. We would also like to thank Fleming’s animal house, flow cytometry (Sofia Grammenoudi), genomics (Vaggelis Harokopos) and microCT facilities. This work was supported by the IMI project BTCure (GA no. 115142-2) and a “Research program for Excellence IKY/Siemens” to GK. We also acknowledge support of this work by the InfrafrontierGR infrastructure co-funded by Greece and the European Union [European Regional Development Fund] under NSRF 2014–2020, MIS 5002135, which provided mouse hosting and phenotyping facilities.

## References

1. Firestein, G. S. Evolving concepts of rheumatoid arthritis. Nature 423, 356–361 (2003).

2. McInnes, I. B., Buckley, C. D. & Isaacs, J. D. Cytokines in rheumatoid arthritis-shaping the immunological landscape. Nat. Rev. Rheumatol. 12, 63–68 (2016).

3. Keffer, J. et al. Transgenic mice expressing human tumour necrosis factor: A predictive genetic model of arthritis. EMBO J. 10, 4025–4031 (1991).

4. Kontoyiannis, D., Pasparakis, M., Pizarro, T. T., Cominelli, F. & Kollias, G. Impaired on/off regulation of TNF biosynthesis in mice lacking TNF AU-rich elements: Implications for joint and gut-associated immunopathologies. Immunity 10, 387–398 (1999).

5. Küsters, S. et al. In vivo evidence for a functional role of both tumor necrosis factor (TNF) receptors and transmembrane TNF in experimental hepatitis. Eur. J. Immunol. 27, 2870–2875 (1997).

6. -Vafeiadou, E. C. et al. Ectopic bone formation and systemic bone loss in a transmembrane TNF-driven model of human spondyloarthritis. 4, 1–14 (2020).

7. Ntari, L. et al. Comorbid TNF-mediated heart valve disease and chronic polyarthritis share common mesenchymal cell-mediated aetiopathogenesis. Ann. Rheum. Dis. 77, 926–934 (2018).

8. Armaka, M. et al. Mesenchymal cell targeting by TNF as a common pathogenic principle in chronic inflammatory joint and intestinal diseases. J. Exp. Med. 205, 331– 337 (2008).

9. Armaka, M., Ospelt, C., Pasparakis, M. & Kollias, G. The p55TNFR-IKK2-Ripk3 axis orchestrates arthritis by regulating death and inflammatory pathways in synovial fibroblasts. Nat. Commun. 9, (2018).

10. Kontoyiannis, D. & Kollias, G. Fibroblast biology. Synovial fibroblasts in rheumatoid arthritis: Leading role or chorus line? Arthritis Res. 2, 342–343 (2000).

11. Dakin, S. G. et al. Pathogenic stromal cells as therapeutic targets in joint inflammation. Nat. Rev. Rheumatol. 14, 714–726 (2018).

12. Neumann, E., Lefèvre, S., Zimmermann, B., Gay, S. & Müller-Ladner, U. Rheumatoid arthritis progression mediated by activated synovial fibroblasts. Trends Mol. Med. 16, 458–468 (2010).

13. Koliaraki, V., Prados, A., Armaka, M. & Kollias, G. The mesenchymal context in inflammation, immunity and cancer. Nat. Immunol. 21, 974–982 (2020).

14. Aidinis, V. et al. Functional analysis of an arthritogenic synovial fibroblast. Arthritis Res. Ther. 5, (2003).

15. Lefèvre, S. et al. Synovial fibroblasts spread rheumatoid arthritis to unaffected joints. Nat. Med. 15, 1414–1420 (2009).

16. Karouzakis, E., Gay, R. E., Gay, S. & Neidhart, M. Epigenetic control in rheumatoid arthritis synovial fibroblasts. Nat. Rev. Rheumatol. 5, 266–272 (2009).

17. Klein, K., Ospelt, C. & Gay, S. Epigenetic contributions in the development of rheumatoid arthritis. Arthritis Res. Ther. 14, (2012).

18. Duroux-Richard, I., Jorgensen, C. & Apparailly, F. What do microRNAs mean for rheumatoid arthritis? Arthritis Rheum. 64, 11–20 (2012).

19. Vicente, R., Noël, D., Pers, Y. M., Apparailly, F. & Jorgensen, C. Deregulation and therapeutic potential of microRNAs in arthritic diseases. Nat. Rev. Rheumatol. 12, 211–220 (2016).

20. Zou, Y. et al. Long noncoding RNA LERFS negatively regulates rheumatoid synovial aggression and proliferation. J. Clin. Invest. 128, 4510–4524 (2018).

21. Jonas, S. & Izaurralde, E. Towards a molecular understanding of microRNA-mediated gene silencing. Nat. Rev. Genet. 16, 421–433 (2015).

22. Bartel, D. P. Metazoan MicroRNAs. Cell 173, 20–51 (2018).

23. Pandis, I. et al. Identification of microRNA-221/222 and microRNA-323-3p association with rheumatoid arthritis via predictions using the human tumour necrosis factor transgenic mouse model. Ann. Rheum. Dis. 71, 1716–1723 (2012).

24. Le Sage, C. et al. Regulation of the p27Kip1 tumor suppressor by miR-221 and miR-222 promotes cancer cell proliferation. EMBO J. 26, 3699–3708 (2007).

25. Garofalo, M. et al. miR-221&222 Regulate TRAIL Resistance and Enhance Tumorigenicity through PTEN and TIMP3 Downregulation. Cancer Cell 16, 498–509 (2009).

26. Pineau, P. et al. miR-221 overexpression contributes to liver tumorigenesis. Proc. Natl. Acad. Sci. U. S. A. 107, 264–269 (2010).

27. Liu, S. et al. A microRNA 221- and 222-mediated feedback loop maintains constitutive activation of NFκB and STAT3 in colorectal cancer cells. Gastroenterology 147, 847-859.e11 (2014).

28. Seeley, J. J. et al. Induction of innate immune memory via microRNA targeting of chromatin remodelling factors. Nature 559, 114–119 (2018).

29. Mikami, Y. et al. MicroRNA-221 and -222 modulate intestinal inflammatory Th17 cell response as negative feedback regulators downstream of interleukin-23. Immunity 54, 514-525.e6 (2021).

30. Abo ElAtta, A. S., Ali, Y. B. M., Bassyouni, I. H. & Talaat, R. M. Upregulation of miR-221/222 expression in rheumatoid arthritis (RA) patients: correlation with disease activity. Clin. Exp. Med. 19, 47–53 (2019).

31. Yang, S. & Yang, Y. Downregulation of microRNAIZl221 decreases migration and invasion in fibroblastIZllike synoviocytes in rheumatoid arthritis. Mol. Med. Rep. 12, 2395–2401 (2015).

32. Galardi, S., Mercatelli, N., Farace, M. G. & Ciafrè, S. A. NF-κkB and c-Jun induce the expression of the oncogenic miR-221 and miR-222 in prostate carcinoma and glioblastoma cells. Nucleic Acids Res. 39, 3892–3902 (2011).

33. Liu, S. et al. A microRNA 221- and 222-mediated feedback loop maintains constitutive activation of NFκB and STAT3 in colorectal cancer cells. Gastroenterology 147, 847-859.e11 (2014).

34. El Gazzar, M. & McCall, C. E. MicroRNAs distinguish translational from transcriptional silencing during endotoxin tolerance. J. Biol. Chem. 285, 20940–20951 (2010).

35. Lodge, R. et al. Host MicroRNAs-221 and -222 Inhibit HIV-1 Entry in Macrophages by Targeting the CD4 Viral Receptor. Cell Rep. 21, 141–153 (2017).

36. Brennan, F. M., Jackson, A., Chantry, D., Maini, R. & Feldmann, M. Inhibitory Effect of Tnfα Antibodies on Synovial Cell Interleukin-1 Production in Rheumatoid Arthritis. Lancet 334, 244–247 (1989).

37. Probert, L., Plows, D., Kontogeorgos, G. & Kollias, G. The type I interleukin-1 receptor acts in series with tumor necrosis factor (TNF) to induce arthritis in TNF-transgenic mice. Eur. J. Immunol. 25, 1794–1797 (1995).

38. Van Den Berg, W. B., Joosten, L. A. B., Kollias, G. & Van De Loo, F. A. J. Role of tumour necrosis factor α in experimental arthritis: Separate activity of interleukin 1β in chronicity and cartilage destruction. Ann. Rheum. Dis. 58, 40–48 (1999).

39. Feldmann, M. Development of anti-TNF therapy for rheumatoid arthritis. Nat. Rev. Immunol. 2, 364–371 (2002).

40. Zwerina, J. et al. Single and Combined Inhibition of Tumor Necrosis Factor, Interleukin-1, and RANKL Pathways in Tumor Necrosis Factor-Induced Arthritis: Effects on Synovial Inflammation, Bone Erosion, and Cartilage Destruction. Arthritis Rheum. 50, 277–290 (2004).

41. Zwerina, J. et al. TNF-induced structural joint damage is mediated by IL-1. Proc. Natl. Acad. Sci. U. S. A. 104, 11742–11747 (2007).

42. Croft, A. P. et al. Distinct fibroblast subsets drive inflammation and damage in arthritis. Nature 570, 246–251 (2019).

43. Clapier, C. R., Iwasa, J., Cairns, B. R. & Peterson, C. L. Mechanisms of action and regulation of ATP-dependent chromatin-remodelling complexes. Nat. Rev. Mol. Cell Biol. 18, 407–422 (2017).

44. Schwenk, F., Baron, U. & Rajewsky, K. A cre-transgenic mouse strain for the ubiquitous deletion of loxP-flanked gene segments including deletion in germ cells. Nucleic Acids Res. 23, 5080–5081 (1995).

45. Armaka, M. et al. Single-cell chromatin and transcriptome dynamics of Synovial Fibroblasts transitioning from homeostasis to pathology in modelled TNF-driven arthritis. bioRxiv 2021.08.27.457747 (2021) doi:10.1101/2021.08.27.457747.

46. Inui, M., Martello, G. & Piccolo, S. MicroRNA control of signal transduction. Nat. Rev. Mol. Cell Biol. 11, 252–263 (2010).

47. Xiao, C. & Rajewsky, K. MicroRNA Control in the Immune System: Basic Principles. Cell 136, 26–36 (2009).

48. O’Connell, R. M., Rao, D. S. & Baltimore, D. MicroRNA regulation of inflammatory responses. Annu. Rev. Immunol. 30, 295–312 (2012).

49. Li, L., Chen, X. P. & Li, Y. J. MicroRNA-146a and human disease. Scand. J. Immunol. 71, 227–231 (2010).

50. Lu, L. F. et al. Function of miR-146a in Controlling Treg Cell-Mediated Regulation of Th1 Responses. Cell 142, 914–929 (2010).

51. Liston, A., Linterman, M. & Lu, L. F. MicroRNA in the adaptive immune system, in sickness and in health. J. Clin. Immunol. 30, 339–346 (2010).

52. Nakasa, T., Shibuya, H., Nagata, Y., Niimoto, T. & Ochi, M. The inhibitory effect of microRNA-146a expression on bone destruction in collagen-induced arthritis. Arthritis Rheum. 63, 1582–1590 (2011).

53. Saferding, V. et al. MicroRNA-146a governs fibroblast activation and joint pathology in arthritis. J. Autoimmun. 82, 74–84 (2017).

54. Stanczyk, J. et al. Altered expression of microRNA in synovial fibroblasts and synovial tissue in rheumatoid arthritis. Arthritis Rheum. 58, 1001–1009 (2008).

55. Blüml, S. et al. Essential role of microRNA-155 in the pathogenesis of autoimmune arthritis in mice. Arthritis Rheum. 63, 1281–1288 (2011).

56. Kurowska-Stolarska, M. et al. MicroRNA-155 as a proinflammatory regulator in clinical and experimental arthritis. Proc. Natl. Acad. Sci. U. S. A. 108, 11193–11198 (2011).

57. Zhu, S. et al. The microRNA miR-23b suppresses IL-17-associated autoimmune inflammation by targeting TAB2, TAB3 and IKK-α. Nat. Med. 18, 1077–1086 (2012).

58. T., P., U., M.-L., R.E., G. & S., G. Fibroblast biology. Role of synovial fibroblasts in the pathogenesis of rheumatoid arthritis. Arthritis Res. 2, 361–367 (2000).

59. Buckley, C. D., Ospelt, C., Gay, S. & Midwood, K. S. Location, location, location: how the tissue microenvironment affects inflammation in RA. Nat. Rev. Rheumatol. 17, 195–212 (2021).

60. Zhang, R. et al. Potential candidate biomarkers associated with osteoarthritis: Evidence from a comprehensive network and pathway analysis. J. Cell. Physiol. 234, 17433–17443 (2019).

61. Rhee, D. K. et al. The secreted glycoprotein lubricin protects cartilage surfaces and inhibits synovial cell overgrowth. J. Clin. Invest. 115, 622–631 (2005).

62. Zhou, Y., Richards, A. M. & Wang, P. MicroRNA-221 Is Cardioprotective and Anti-fibrotic in a Rat Model of Myocardial Infarction. Mol. Ther. - Nucleic Acids 17, 185– 197 (2019).

63. Ding, L. et al. Circular RNA circ-DONSON facilitates gastric cancer growth and invasion via NURF complex dependent activation of transcription factor SOX4. Mol. Cancer 18, 1–11 (2019).

64. Liu, T. et al. The transcription factor Zfp90 regulates the self-renewal and differentiation of hematopoietic stem cells article. Cell Death Dis. 9, (2018).

65. Lignitto, L. et al. Nrf2 Activation Promotes Lung Cancer Metastasis by Inhibiting the Degradation of Bach1. Cell 178, 316-329.e18 (2019).

66. Rothe, J. et al. Mice lacking the tumour necrosis factor receptor 1 are resistant to IMF-mediated toxicity but highly susceptible to infection by Listeria monocytogenes. Nature 364, 798–802 (1993).

67. Bouxsein, M. L. et al. Guidelines for assessment of bone microstructure in rodents using micro-computed tomography. J. Bone Miner. Res. 25, 1468–1486 (2010).

68. Armaka, M., Gkretsi, V., Kontoyiannis, D. & Kollias, G. A standardized protocol for the isolation and culture of normal and arthritogenic murine synovial fibroblasts. Protoc. Exch. 1–10 (2009) doi:10.1038/nprot.2009.102.

69. Granja, J. M. et al. Author Correction: ArchR is a scalable software package for integrative single-cell chromatin accessibility analysis (Nature Genetics, (2021), 53, 3, (403-411), 10.1038/s41588-021-00790-6). Nat. Genet. 53, 935 (2021).

70. Corces, M. R. et al. The chromatin accessibility landscape of primary human cancers. Science (80-.). 362, (2018).

71. Schep, A. N., Wu, B., Buenrostro, J. D. & Greenleaf, W. J. ChromVAR: Inferring transcription-factor-associated accessibility from single-cell epigenomic data. Nat. Methods 14, 975–978 (2017).

