## Supplementary Figures for "miR-221/222 drive synovial fibroblast expansion and pathogenesis of TNF-mediated arthritis"

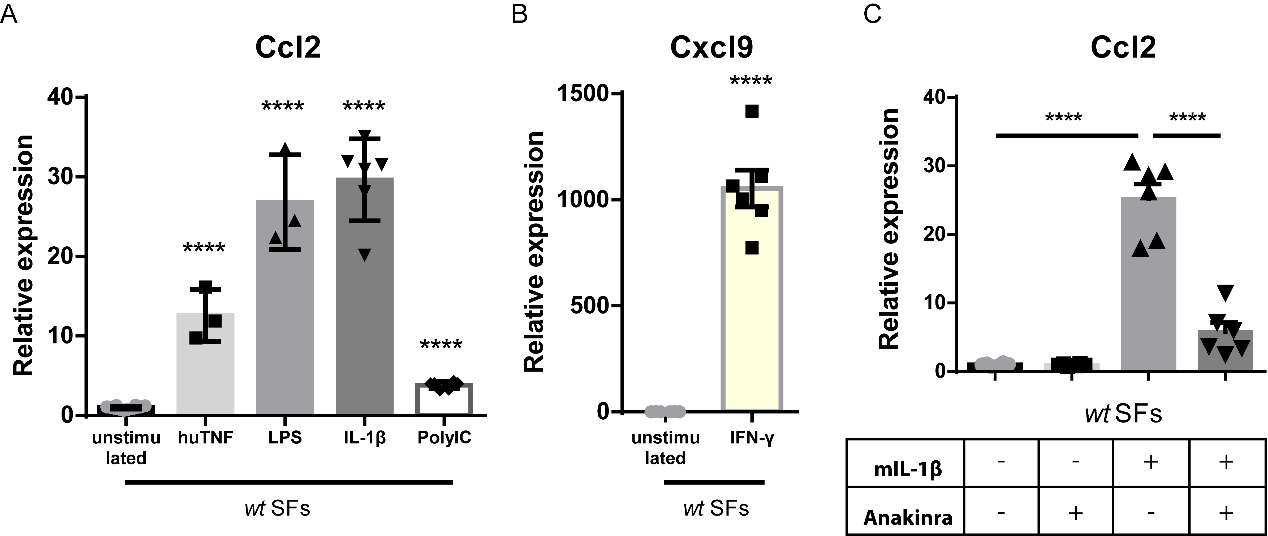

Figure S1: **SFs respond to inflammatory signals**.

A) qRT-PCR analysis of Ccl2 expression in cultured WT SFs after 24h stimulation with TNF, LPS, IL1-β and PolyIC (n = 3-6). WT unstimulated SFs served as reference.

B) qRT-PCR analysis of Cxcl9 expression in cultured WT SFs after 24h stimulation with IFN-γ (n = 6). WT unstimulated SFs served as reference for normalization.

C) qRT-PCR analysis of Ccl2 expression in cultured WT SFs after 24h stimulation with TNF and IL1-β in the presence or absence of anakinra (n = 3-6). Expression was normalized to the levels detected in WT unstimulated SFs.

In all expression analysis experiments B2m was used as a housekeeping gene for normalization.

Data represent mean ± SEM. *p < 0.05, **p < 0.01, ***p < 0.001, ****p < 0,0001, ns = not significant.

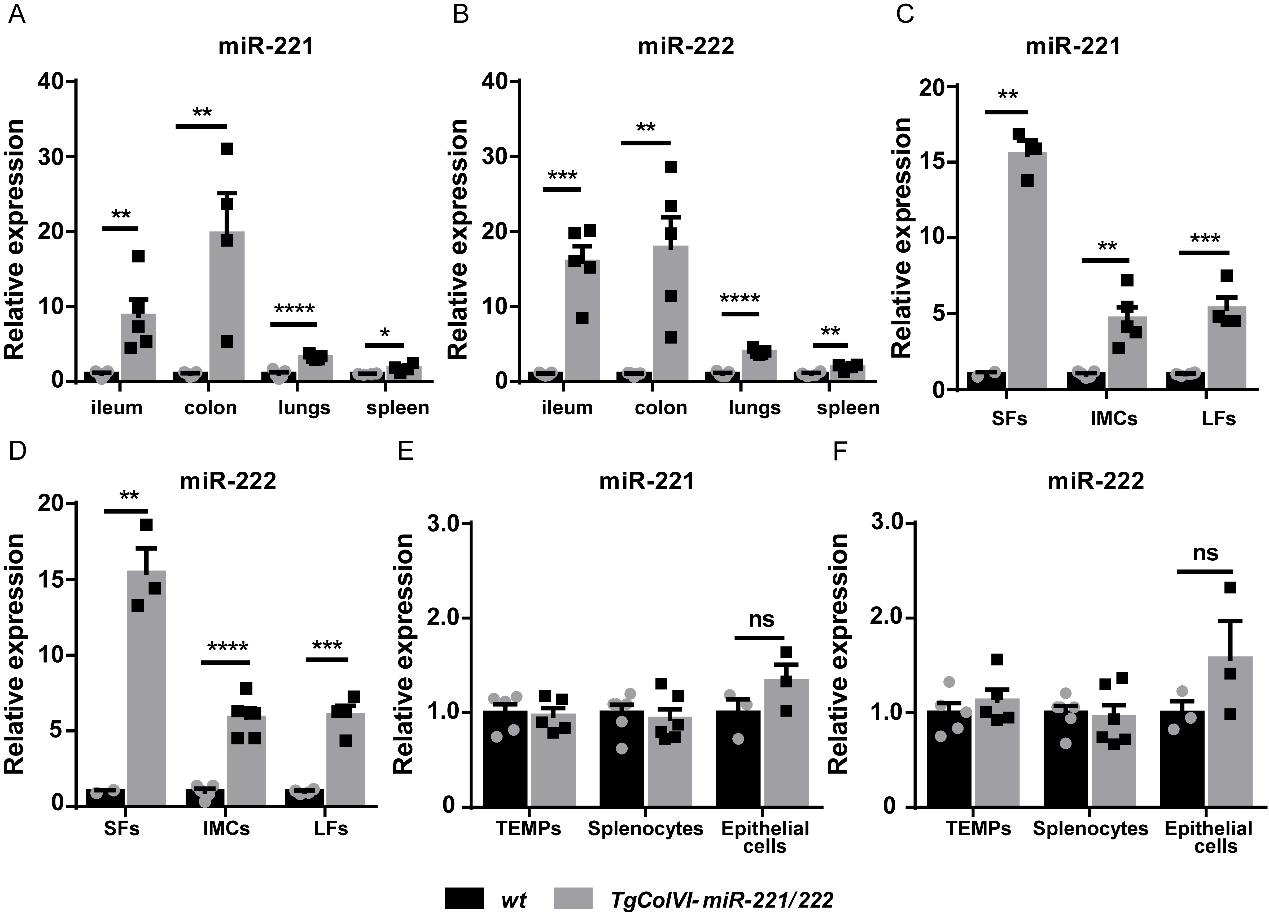

Figure S2: ***TgColVI-miR-221/222* mice target tissues and cells of mesenchymal origin**.

Expression levels of miR-221 and miR-222 as determined by qRT-PCR from

A-B) Ileum, colon, lung and spleen from *TgColVI-miR-221/222* mice.

C-D) SFs, intestinal mesenchymal cells (IMCs) and lung fibroblasts (LFs) from *TgColVI-miR-221/222* mice.

E-F) Peritoneal macrophages (TEMPs), splenocytes and epithelial cells from *TgColVI-miR-221/222* mice.

Expression levels were normalized to the levels seen in WT mice (n = 3- 6). u6 was used for normalization.

Data represent mean ± SEM. *p < 0.05, **p < 0.01, ***p < 0.001, ****p < 0,0001, ns = not significant.
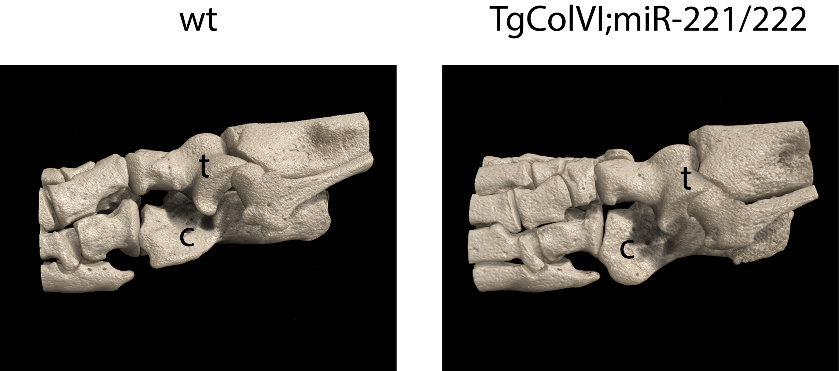

Figure S3: **miR-221/222 overexpression may affect bone physiology**.

Representative microCT images from WT and *TgColVI-miR-221/222* ankle joint area. t: talus, c: calcaneous.

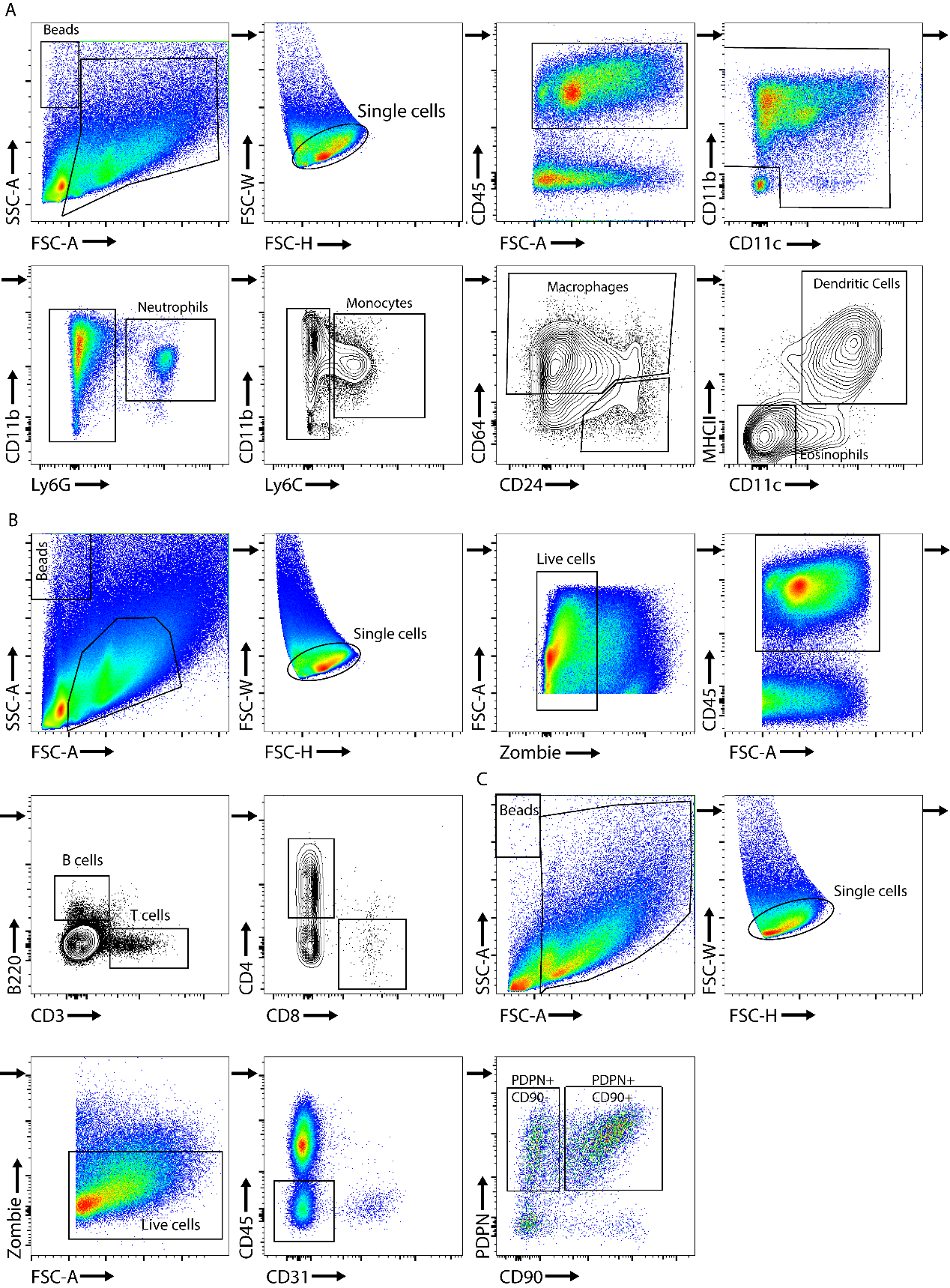

Figure S4: **FACS gating strategies.** (A) FACS gating strategy for myeloid cell infiltration in the ankle joints. The cell suspension was gated for live CD45^+^. In the CD45^+^ population, we characterized the abundance of macrophages, monocytes, neutrophils, dendritic cells and eosinophils.

B) FACS gating strategy for lymphocyte cell infiltration in the ankle joints. The cell suspension was gated for live CD45^+^ and we plotted for CD4^+^ and CD8^+^ T cells.

C) FACS gating strategy for synovial fibroblast populations in the ankle joints. The cell suspension was gated for live CD45^-^CD31^-^ and we gated for PDPN and CD90 expression on fibroblasts.

Single-cell suspensions from ankle joints were gated for live cells using zombie green or NIR. Counting beads were used for quantification of different cell subpopulations.

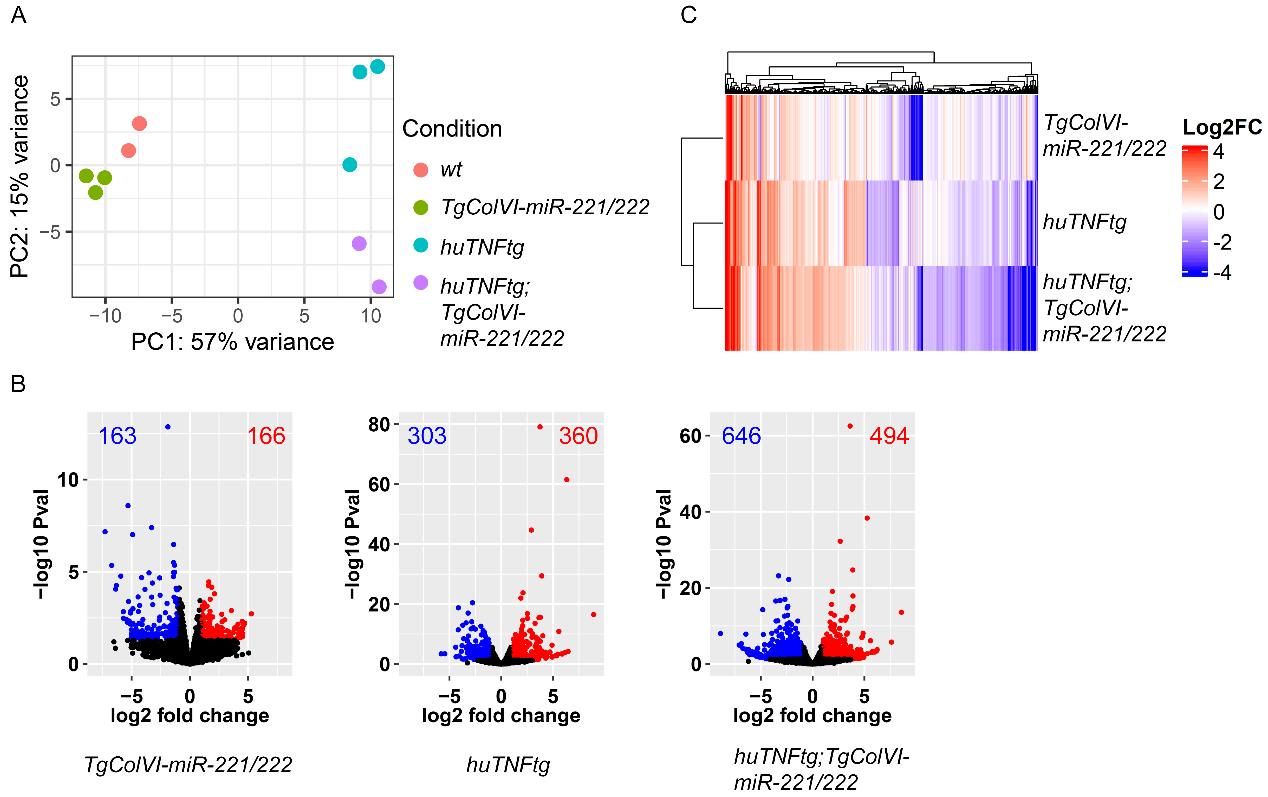

Figure S5: **Comparisons between samples used for bulk RNA sequencing**.

A) Principal Component Analysis (PCA plot) of cultured SFs from 8 week-old WT (n = 2), *TgColVI-miR-221/222* (n = 3), *huTNFtg* (n = 3) and *huTNFtg;TgColVI-miR-221/222* mice (n = 2) mice.

B) Volcano plots of deregulated genes in SFs from *TgColVI-miR-221/222*, *huTNFtg* and *huTNFtg;TgColVI-miR-221/222* compared to WT SF expression profile. Not significantly deregulated genes are depicted with black color, significantly upregulated genes (p-value < 0.05, log2FC > 1) with red and significantly downregulated genes (p-value < 0.05, log2FC < -1) with blue.

C) Heatmap showing log2 fold change values for the deregulated genes and contrasts plotted in B. Hierarchical clustering has been performed for both genes and samples.

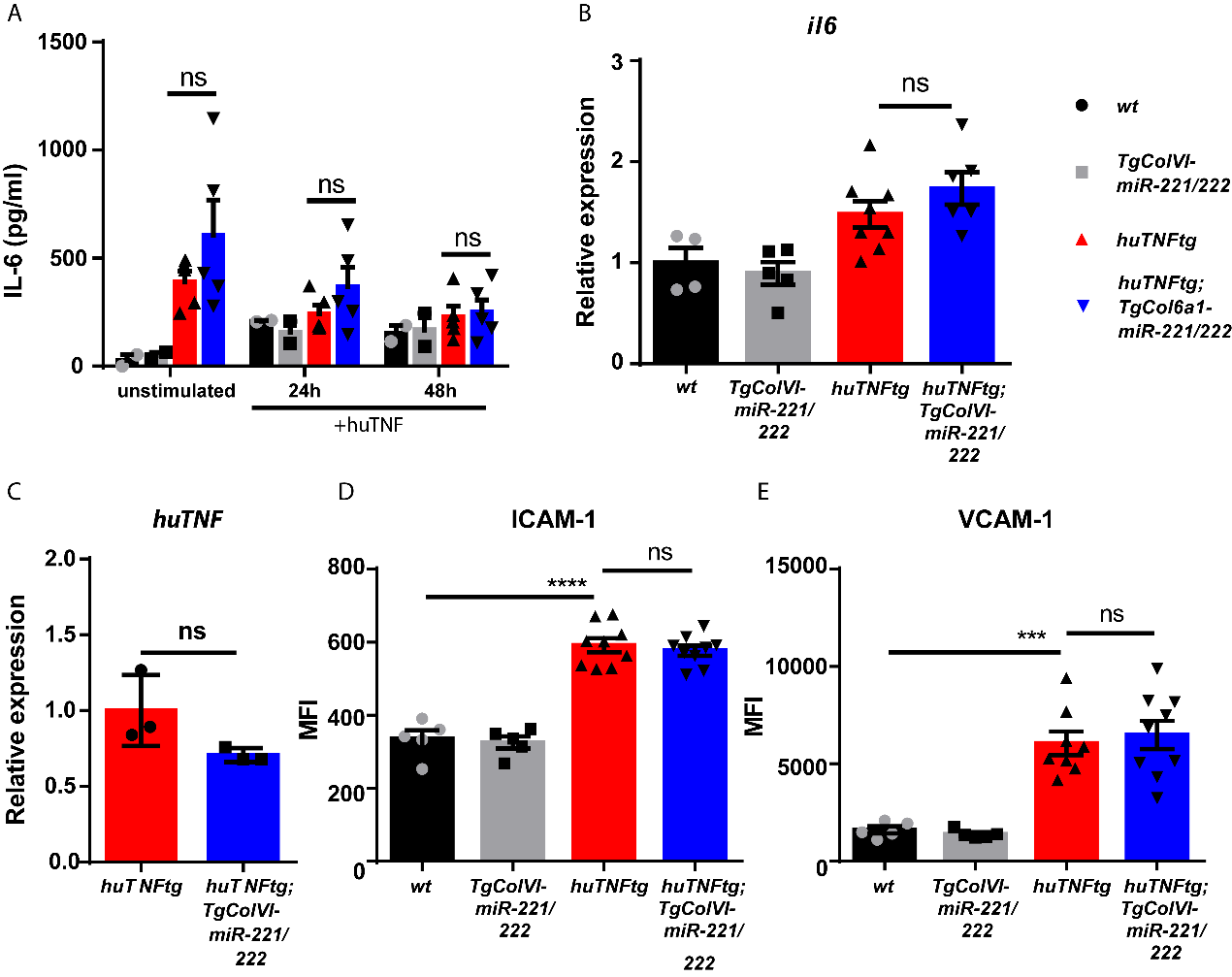

Figure S6: **miR-221/222 do not regulate inflammatory signaling in SFs**. A) IL-6 protein levels measured by ELISA in supernatants from WT (n = 2), *TgColVI-miR-221/222* (n = 2), *huTNFtg* (n = 5) and *huTNFtg;TgColVI-miR-221/222* mice (n = 5) SFs unstimulated or TNF stimulated for 24h or 48h.

B) qRT-PCR analysis of I*l6* from WT (n = 4), *TgColVI-miR-221/222* (n = 5), *huTNFtg* (n = 8) and *huTNFtg;TgColVI-miR-221/222* mice (n = 6) SFs. WT levels were used for normalization. B2m was used as a housekeeping gene.

C) qRT-PCR analysis of *huTNF* in SFs from huTNFtg (n=3) and huTNFtg;TgColVI-miR-221/222 mice (n=3). *huTNFtg* samples were used for normalization. B2m was used as a housekeeping gene.

D-E) Flow cytometry analysis of ICAM-1 and VCAM-1 expression as MFI in SFs from WT (n = 4), *TgColVI-miR-221/222* (n = 5), *huTNFtg* (n = 8-9) and *huTNFtg;TgColVI-miR-221/222* mice, (n = 9).

Data represent mean ± SEM. *p < 0.05, **p < 0.01, ***p < 0.001, ****p < 0,0001, ns = not significant.

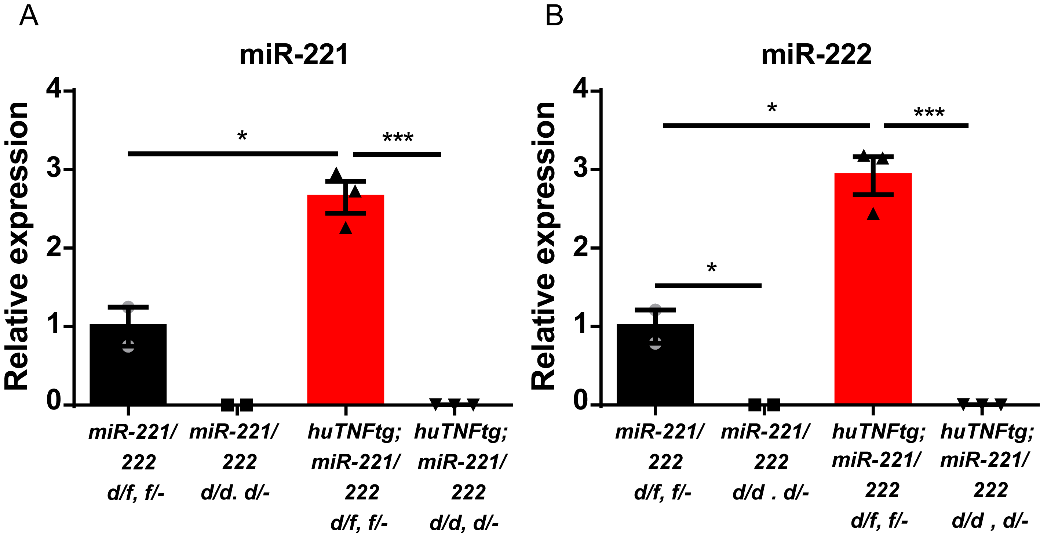

Figure S7: **SFs from miR-221/222 -/- lack miR-221/222 expression.**

A-B) qRT-PCR for miR-221 and -222 levels in cultured SFs from 9 week-old *miR-221/222 d/f, miR-221/222 -/-, huTNFtg;miR-221/222 d/f* and *huTNFtg;miR-221/222 -/-* (n = 2-3). Expression of miR-221/222 d/f SFs was used for normalization.

In all experiments *u6* was used as a housekeeping gene for normalization.

Data represent mean ± SEM. *p < 0.05, **p < 0.01, ***p < 0.001, ****p < 0,0001, ns = not significant.

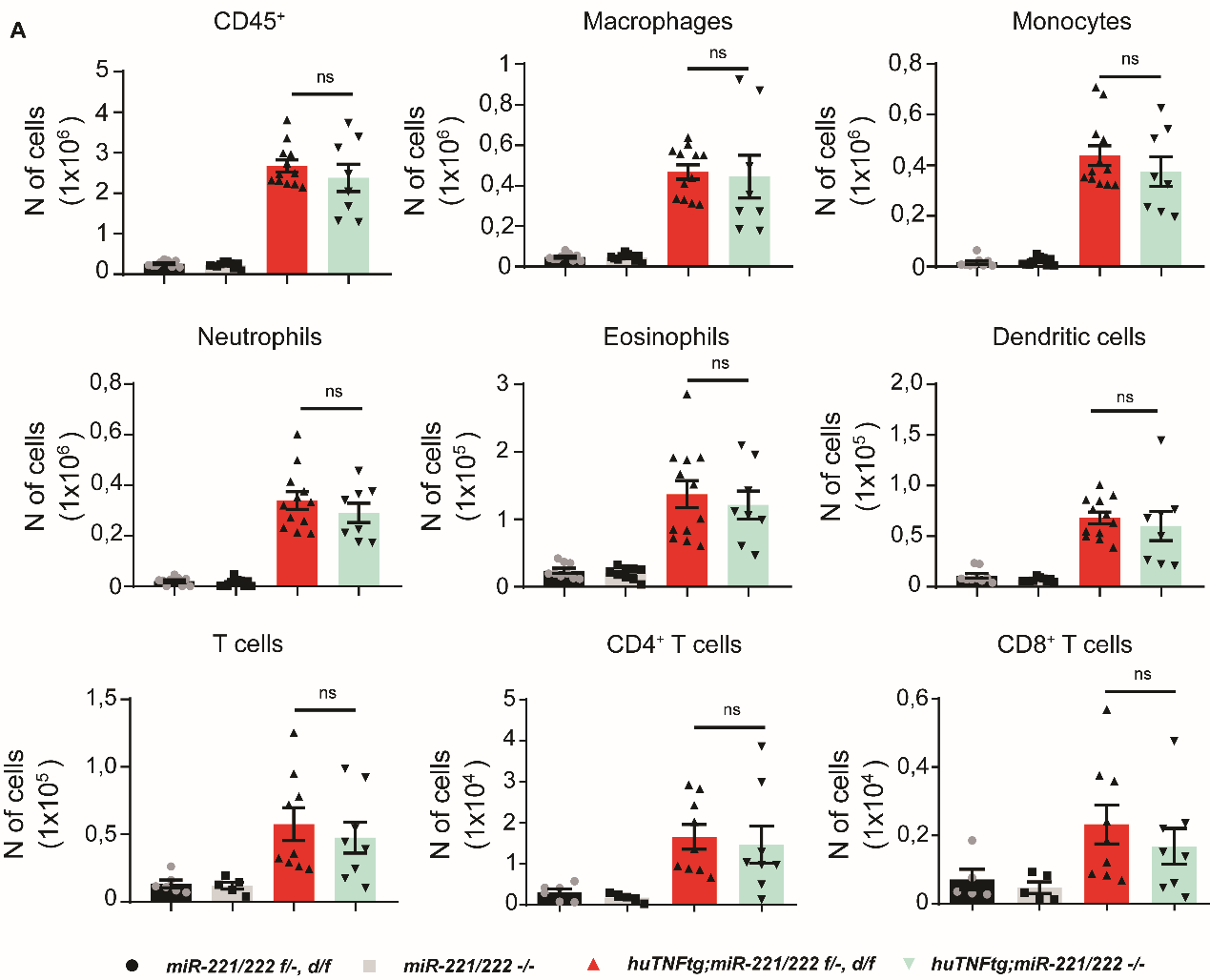

Figure S8: **miR-221/222 -/- deletion in experimental arthritis does not regulate inflammatory influx in the joints.**

Infiltration of CD45^+^ cells, macrophages, monocytes, neutrophils, eosinophils, dendritic cells, CD4^+^ T cells and CD8^+^ T cells in the ankle joints of 9 week-old *miR-221/222 f/-, d/f* (n = 6-9), *miR-221/222 -/-* (n = 5-10), *huTNFtg;miR-221/222 f/-, d/f* (n = 9-12) *huTNFtg;miR-221/222 -/-* mice (n = 8-10) by FACS analysis (from three independent experiments).

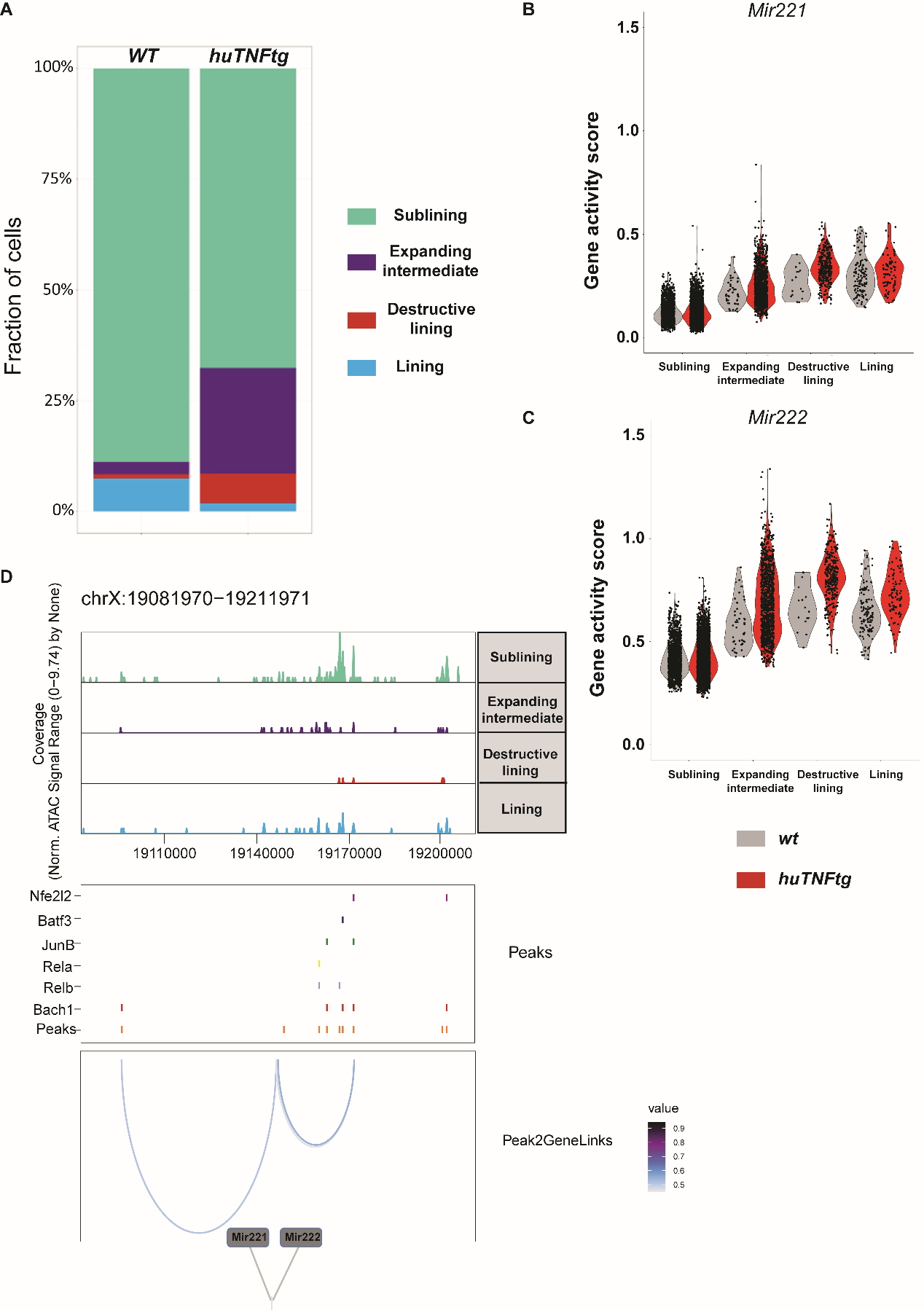

Figure S9: **miR-221/222 gene activity in SF clusters in normal and arthritic state.**

A) Stacked barchart showing relative abundances (%) of cell clusters (as described in Figure 6A) across *WT* and *huTNFtg* samples. Both intermediate and destructive lining populations are expanding in disease state.

B-C) Violin plots exhibiting gene activity scores for Mir221 and Mir222 genes. Split view is selected, in order to highlight the differences between *WT (shown in grey)* and *huTNFtg (shown in red)* samples across the different clusters.

D) Genome accessibility track visualization of the extended regulatory space of Mir221 and Mir222 (chrX:19,081,970-19,211,971), with TF binding site and peak-to-gene linkage information, in *WT* sample. Upper, the genome track shows increased accessibility in intermediate and destructive lining clusters. Middle, all reproducible peaks are shown, coupled with annotated CISBP binding information for Bach1, Rela, Relb, JunB, Batf3 and Nfe2l2 (Nrf2) TFs. Lower, putative regulatory linkages between Mir221-Mir222 genes and reproducible peaks are illustrated. Links between genes and peaks are colored by correlation (Pearson coefficient) of peak accessibility and gene activity scores.
